# Mitochondrial Phosphatidylethanolamine Directly Regulates UCP1 to Promote Brown Adipose Thermogenesis

**DOI:** 10.1101/2022.08.26.505474

**Authors:** Jordan M. Johnson, Alek D. Peterlin, Enrique Balderas, Elahu G. Sustarsic, J. Alan Maschek, Marisa J. Lang, Alejandro Jara-Ramos, Vanja Panic, Jeffrey T. Morgan, Claudio J. Villanueva, Alejandro Sanchez, Jared Rutter, Irfan J. Lodhi, James E. Cox, Kelsey H. Fisher-Wellman, Dipayan Chaudhuri, Zachary Gerhart-Hines, Katsuhiko Funai

## Abstract

Thermogenesis by uncoupling protein 1 (UCP1) is one of the primary mechanisms by which brown adipose tissue (BAT) increases energy expenditure. UCP1 resides in the inner mitochondrial membrane (IMM), where it dissipates membrane potential independent of ATP synthase. Here we provide evidence that mitochondrial phosphatidylethanolamine (PE) directly regulates UCP1-dependent proton conductance across IMM to modulate thermogenesis. Mitochondrial lipidomic analyses revealed PE as a signature molecule whose abundance bidirectionally responds to changes in thermogenic burden. Reduction in mitochondrial PE by deletion of phosphatidylserine decarboxylase (PSD) made mice cold intolerant and insensitive to β3 adrenergic receptor agonist-induced increase in whole-body oxygen consumption. High-resolution respirometry and fluorometry of BAT mitochondria showed that loss of mitochondrial PE specifically lowers UCP1-dependent respiration without compromising electron transfer efficiency or ATP synthesis. These findings were confirmed by a reduction in UCP1 proton current in PSD-deficient mitoplasts. Thus, PE performs a previously unknown role as a temperature-responsive rheostat that regulates UCP1-dependent thermogenesis.

## INTRODUCTION

Nearly two thirds of adults in the United States are overweight or obese, putting them at high-risk to develop cardiovascular disease, type 2 diabetes, and cancer (Hales et al., 2020; Schelbert, 2009). Increasing thermogenesis in adipose tissue is a potential means of increasing energy expenditure and preventing hyperglycemia. Uncoupling protein 1 (UCP1), an integral membrane protein that resides in the inner mitochondrial membrane (IMM) is largely responsible for thermogenesis in brown and beige adipocytes. UCP1 promotes inefficient oxidative phosphorylation (OXPHOS) by dissipating the IMM proton gradient independent of ATP synthesis. Overexpression of UCP1 in adipose tissue is sufficient to prevent diet-induced obesity (Kopecký et al., 1996).

In both mice and humans, UCP1-positive brown and/or beige adipocytes are primarily activated by changes in environmental temperature. While transcriptional control of UCP1 has been well described, relatively less is known about mechanisms that regulate UCP1 protein function. The IMM is a protein-rich phospholipid bilayer, whose lipid composition can substantially impact enzyme activity (Cogliati et al., 2016). Cone-shaped phosphatidylethanolamine (PE) and cardiolipin (CL) are particularly enriched in cristae, where they are thought to create the negative curvature and support the function of the complexes of the electron transport system (Heden et al., 2016; Mejia and Hatch, 2016; Saks et al., 2003). Mutations in the genes of mitochondrial PE or CL biosynthesis are known to cause severe mitochondrial disease, characterized by poor cristae formation, impaired capacity for OXPHOS, blunted efficiency of electron transfer, reduced electron transport system (ETS) complex activity, and decreased supercomplex assembly (Barth et al., 1983; Funai et al., 2020; Girisha et al., 2019; Powers et al., 2013; Tasseva et al., 2013; Zhao et al., 2019). CL and PE abundance both increase with cold exposure, and CL is required for thermogenesis partly via the transcriptional regulation of UCP1 (Marcher et al., 2015; Sustarsic et al., 2018). However, it is unknown whether these lipids directly regulate UCP1 activity to modulate thermogenesis.

Here we sought to determine whether interventions that activate BAT thermogenesis were associated with distinct phospholipid signatures in mitochondria. Comprehensive mitochondrial lipidomic analyses of BAT suggested that PE, not CL, is the primary lipid class responsive to altering energy demand in BAT. Genetic ablation of mitochondrial PE biosynthesis, but not CL, reduced UCP1-dependent respiration. Further, direct patch-clamp measurements of mitoplasts synthesized from these mitochondria showed that reduction in PE lowered UCP1 proton current. These findings implicate mitochondrial PE as an energy-responsive IMM element that regulates UCP1 activity and BAT thermogenesis.

## RESULTS

### PE is the energy-responsive phospholipid in BAT mitochondria

Interventions that influence energy balance such as diet, exercise, and ambient temperature are known to alter the mitochondrial lipid composition in a variety of tissues (Chung et al., 2018; Mendham et al., 2021; O’Shea et al., 2009; Ostojic et al., 2013; Stanley et al., 2012). We set out to determine whether interventions that alter the energic burden of BAT thermogenesis coincide with changes in the mitochondrial lipid milieu (Figure 1A). We first compared wildtype C57BL/6J mice that were housed at 6.5 °C (Cold) for 1 week to 22 °C (RT)-housed controls. Deep phenotyping of BAT mitochondria from these mice indicated markedly elevated respiration (Figure S1A) with a modest reduction in the rate of ATP synthesis in the cold-housed mice (Figure S1B). The reduced efficiency for OXPHOS (ATP/O, Figure S1C) was not due to increased electron leak (Figure S1D) but because of a robust increase in UCP1-dependent respiration (Figure 1B, quantified by subtracting *J*O_2_ with UCP1 inhibition from total *J*O_2_). Consistent with these findings, the increase in mitochondrial respiration occurred concomitant with increased abundance of UCP1 in mitochondria, without changes in the quantity of ETS enzymes (Figure 1C).

**Figure 1:**
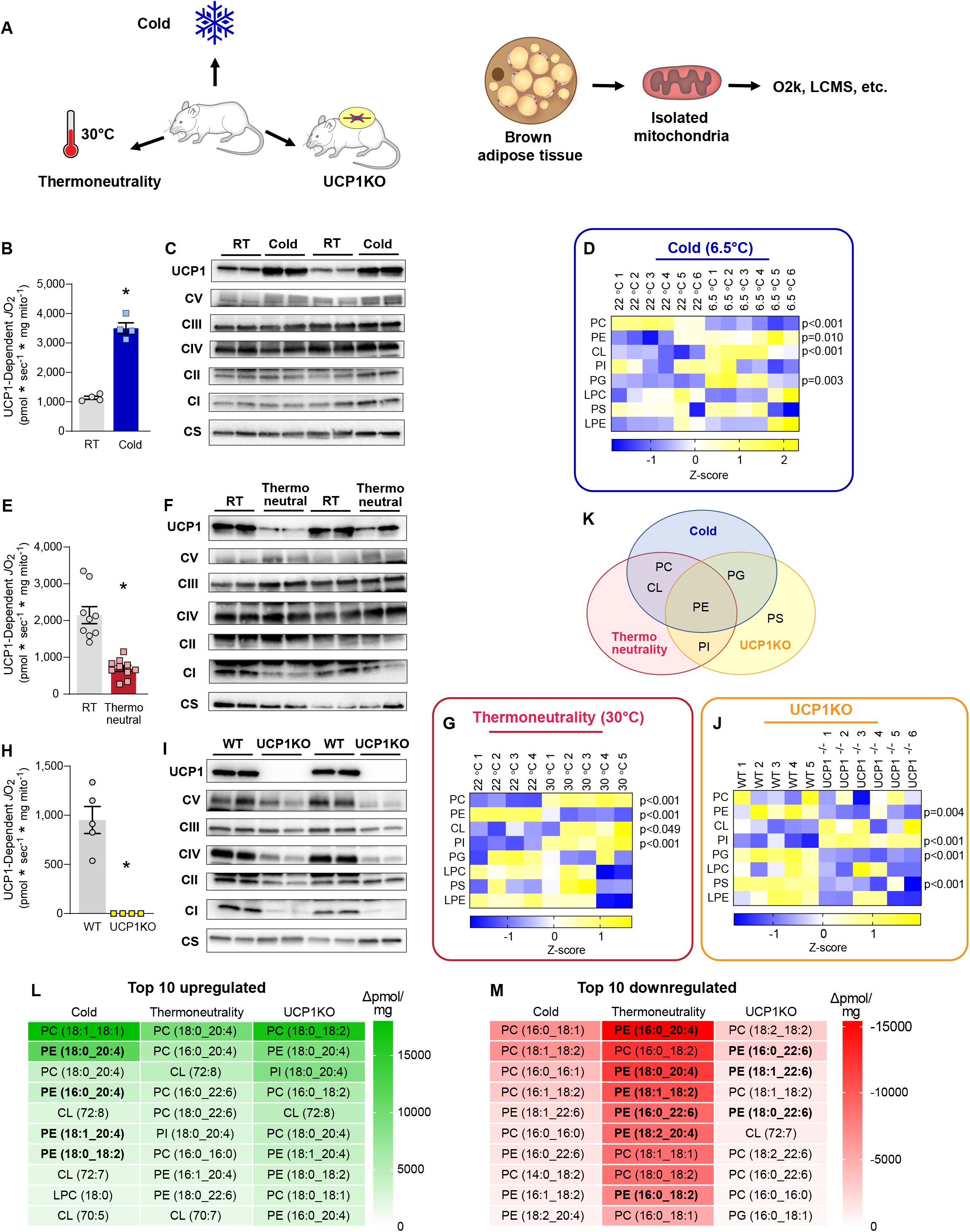
Brown adipose mitochondrial lipidome is highly responsive to changes in thermogenic burden. (A) Experimental design to assess the influences of cold, thermoneutrality, or UCP1 knockout on BAT mitochondrial energetics and lipidome. (B) UCP1-dependent respiration in BAT mitochondria from C57BL/6J mice housed at RT or 6.5 °C for 7 days. n=4/group. (C) Protein abundance of UCP1, ETS subunits, and citrate synthase in BAT mitochondria isolated from C57BL/6J mice housed at RT or 6.5 °C for 7 days. (D) A summary heatmap of changes in BAT mitochondrial phospholipidome in C57BL/6J mice housed at RT or 6.5 °C for 7 days. Abundance of lipids in each lipid class was derived from individual lipid species in Figure S1. n=6/group. (E) UCP1-dependent respiration stimulated in BAT mitochondria from C57BL/6J mice housed at RT or 30 °C for 30 days. n=8/group. (F) Protein abundance of UCP1, ETS subunits, and citrate synthase in BAT mitochondria from C57BL/6J mice housed at RT or 30 °C for 30 days. (G) A summary heatmap of changes in BAT mitochondrial phospholipidome in C57BL/6J mice housed at RT or 30 °C for 30 days. Abundance of lipids in each lipid class was derived from individual lipid species in Figure S2. n=4-5/group. (H) UCP1-dependent respiration in BAT mitochondria from WT and UCP1KO mice. n=8/group. (I) Protein abundance of UCP1, ETS subunits, and citrate synthase in BAT mitochondria from WT and UCP1KO mice. (J) A summary heatmap of changes in BAT mitochondrial phospholipidome in WT and UCP1KO mice, n=5-6/group. (K) Venn Diagram demonstrating PE as the only class of lipids that is influenced by cold, thermoneutrality, and UCP1 knockout. (L) A list of top 10 mitochondrial lipid species that are upregulated with cold, thermoneutrality, or UCP1 knockout. (M) A list of top 10 mitochondrial lipid species that are downregulated with cold, thermoneutrality, or UCP1 knockout. Data are presented as ± S.E.M. *Denotes of p-value of 0.05 or less.

Comprehensive lipid mass spectrometric analyses of BAT mitochondria revealed significant increases in PE, CL, and phosphatidylglycerol (PG), and a decrease in phosphatidylcholine (PC) (Figure 1D, S1E-L). Cold-stimulated increase in CL and PG were consistent with our previous report (Sustarsic et al., 2018). We also observed an equally robust increase in mitochondrial PE, analogous to exercise-induced increase in mitochondrial PE that was observed in skeletal muscle (Heden et al., 2019). The changes in these lipids coincided with increased expression of the biosynthetic enzymes responsible for mitochondrial PE and CL production (Figure S1M).

Next, we examined BAT mitochondria from wildtype C57BL/6J mice that were housed at 30 °C (Thermoneutrality) or 22 °C (RT) for 4 weeks. Mice exhibit a modestly-elevated BAT-mediated thermogenesis at room temperature (∼22° C) compared to thermoneutrality (Gordon, 2012; Speakman and Keijer, 2013). As expected, mitochondrial phenotyping experiments showed reduced respiration (Figure S2A) and increased capacity for ATP synthesis in the thermoneutral-housed mice compared to RT-housed mice (Figure S2B). The increased efficiency of OXPHOS under thermoneutral conditions (Figure S2C) was not due to altered electron leak (Figure S2D) but because of markedly diminished UCP1-dependent respiration (Figure 1E). Reduced UCP1 content in thermoneutral mitochondria was consistent with the diminished UCP1-dependent *J*O_2_ (Figure 1F).

Thermoneutrality influenced the BAT mitochondrial lipidome in some unexpected ways (Figure 1G and S2E-M). Abundance of CL was higher, not lower, in mice housed at thermoneutrality compared to mice housed at RT. Instead, PE was the only lipid class whose abundance was downregulated with thermoneutrality. In addition to CL, PC and phosphatidylinositol (PI) were also increased with thermoneutrality.

To further show that PE, not CL, is the energy-responsive lipid in the IMM, we next examined UCP1 null (UCP1KO) mice. BAT mitochondria from UCP1KO mice are similar to mitochondria from thermoneutral conditions in that they lack UCP1-driven thermogenesis. Indeed, UCP1 deletion reduced respiration (Figure S3A) and increased efficiency of OXPHOS (Figure S3B&C) that was due to a lack of UCP1-dependent respiration (Figure 1H&I and S3D). Phospholipidomic analyses revealed changes in the mitochondrial lipid milieu with UCP1 deletion that were remarkably similar to what was observed in changes with thermoneutrality (Figure 1J and S3E-M). CL was not responsive to UCP1 deletion (trending for an increase rather than a decrease). Instead, PE, PG, and PS were lower while phosphatidylinositol (PI) was higher in UCP1KO mice compared to wildtype controls. Together, PE was the only energy responsive lipids across the three models that were examined (Figure 1K). PE is highly abundant (40% of total BAT mitochondrial phospholipids) and is important for OXPHOS (Heden et al., 2019; Tasseva et al., 2013; van der Veen et al., 2014). Indeed, PE ranked consistently high among the lipid species whose abundance was upregulated with cold and downregulated with thermoneutrality or UCP1 knockout when compared to relevant control samples (Figure 1L&M).

Based on these observations, we hypothesized that PE was a likely candidate to regulate UCP1 function to modulate thermogenesis. Nevertheless, based on our previous findings that CL is an important regulator of BAT thermogenesis, we further examined the role of both PE and CL in UCP1-dependent thermogenesis.

### BAT-specific deletion of CL synthase impairs thermogenesis

We previously demonstrated that deletion of CL synthase (CLS) (Figure 2A) driven by adiponectin-Cre, tamoxifen-inducible Rosa26-Cre, and tamoxifen-inducible UCP1-Cre (UCP1 Cre-ERT2) impairs thermogenesis and leads to insulin resistance (Sustarsic et al., 2018). The majority of these experiments were performed in adiponectin-Cre driven CLS knockout mice. Here we performed additional studies on CLS knockout driven by the inducible UCP1-Cre (CLS-iBKO) to conduct a more in-depth mitochondrial phenotyping.

**Figure 2:**
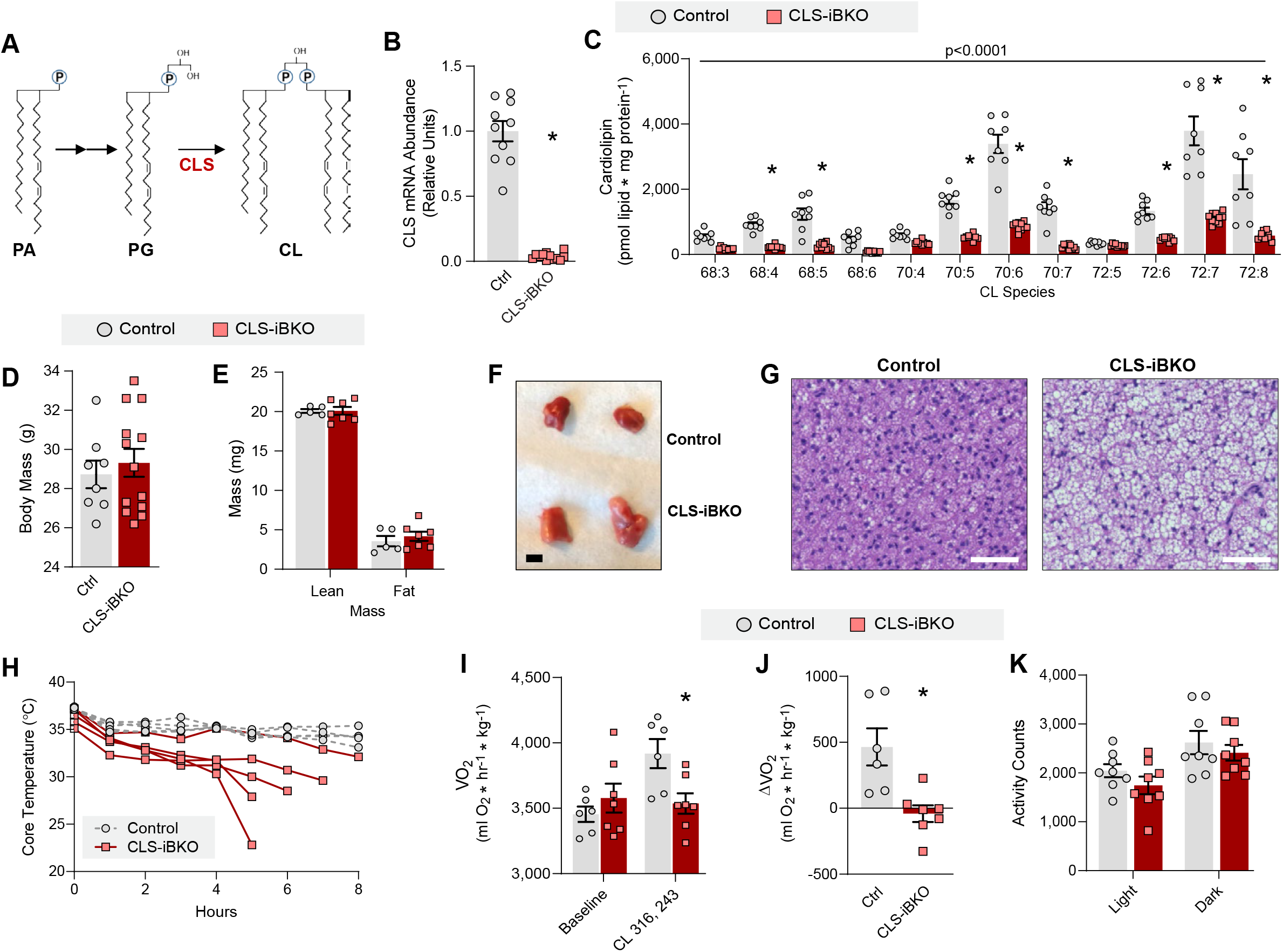
Loss of mitochondrial CL impairs brown adipose thermogenesis. (A) A schematic for mitochondrial CL biosynthesis. (B) CLS mRNA abundance in BAT from control and CLS-iBKO mice. n=10-12/group. (C) Abundance of mitochondrial CL species in BAT from control and CLS-iBKO mice. n=3-8/group. (D) Body mass. n=8-13/group. (E) Body composition. n=5-7/group. (F) Representative intrascapular BAT depots from control and CLS-iBKO mice. (G) Hematoxylin and eosin staining. Scale bar=50 µM. (H) Core body temperature of mice subjected to an acute cold-tolerance test at 4° C. n=5/group. (I) Whole-body oxygen consumption in mice before and after injection of CL-316,243. n=6-7/group. (J) Changes in whole-body oxygen consumption following administration of CL-316,243. n=6-7/group. (K) Spontaneous movement in metabolic cages. n=8/group. Data are presented as ± S.E.M. *Denotes of p-value of 0.05 or less.

As expected, the CLS mRNA abundance was reduced in CLS-iBKO mice compared to control mice (Figure 2B). However, due to its relatively slow turnover, CL levels were not substantially reduced until 4-weeks post-tamoxifen injection (Figure 2C). Thus, we performed all subsequent analyses of the CLS-iBKO and control mice at 4-weeks post-tamoxifen injection. Consistent with CLS’s activity of synthesizing CL from PG, CLS deletion also increased mitochondrial PG compared to controls (Figure S4A). CLS-iBKO and control mice did not differ in body mass, lean mass, or fat mass (Figure 2D&E), but BAT was both larger and paler in CLS-iBKO mice (Figure 2F, S4B). Histological sections of BAT revealed increased lipid accumulation in the CL deficient samples (Figure 2G), potentially suggesting impaired thermogenic capacity. Indeed, reduced CL levels impaired cold tolerance when mice were challenged at 4° C (Figure 2H). Furthermore, although administration of the β3-agonist CL-316,243 increased VO_2_ in control mice, CLS-iBKO mice were refractory to CL-316,243-induced changes in VO_2_, which indicates defective thermogenesis (Figure 2I&J). 24-hr calorimetry at room temperature showed no effect of CLS deletion on VO_2_, RER, and activity, suggesting that these animals are otherwise not hypoenergetic (Figure 2K and S4B&C).

### CLS deletion impairs thermogenesis independent of its effect on UCP1-dependent respiration

Based on CL’s localization to the IMM and its effect on OXPHOS, we hypothesized that CLS deletion lowers CL to reduce UCP1 activity. To test this, we phenotyped BAT mitochondria from control and CLS-iBKO mice. Electron micrograph of BAT surprisingly revealed no substantial deformities in mitochondrial shape or cristae density in CLS-iBKO derived samples compared to wildtype-derived samples (Figure 3A). Indeed, quantification of mitochondrial density by mtDNA/nucDNA (Figure 3B), protein quantifications of UCP1, complex I-V, and citrate synthase (Figure 3C), and citrate synthase activity (Figure 3D) showed no difference between BAT from control and CLS-iBKO mice. Bioenergetic experiments further revealed unexpected effects of CLS deletion on mitochondria. ADP-dependent respiration was elevated, not reduced, in CLS-lacking mitochondria (Figure 3E). The increased respiration coincided with an increased rate of ATP synthesis (Figure 3F) and no overall effect on the ATP/O ratio (Figure 3G) or electron leak (Figure S4D). Strikingly, in stark contrast to our hypothesis, CLS deletion had no effect on UCP1-dependent respiration (Figure 3H).

**Figure 3:**
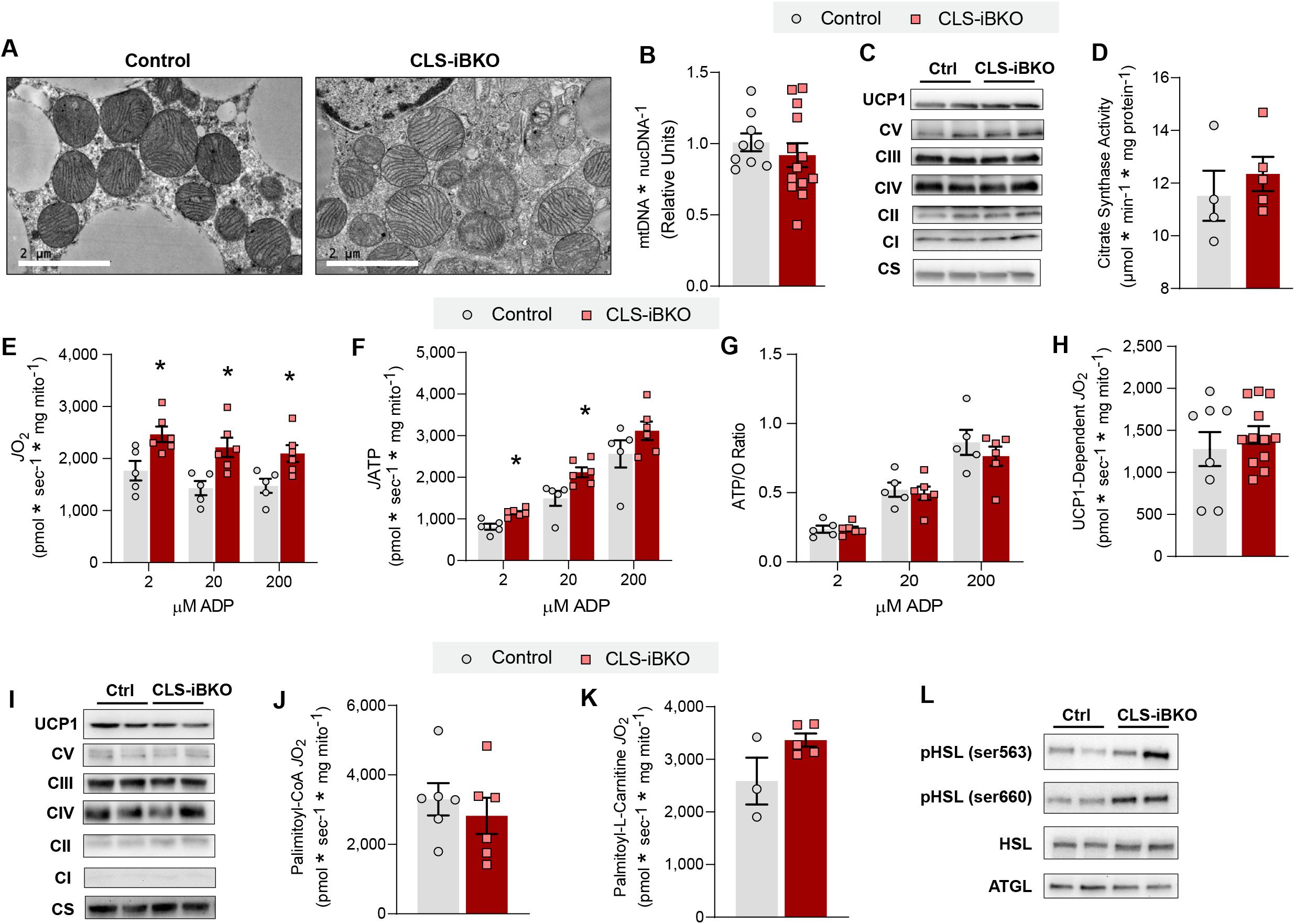
CL deficiency does not impair UCP1-dependent respiration. (A) Transmission electron microscopy images of BAT mitochondria from control and CLS-iBKO mice. Scale bar=2 µM. (B) Mitochondrial DNA levels (mtDNA) normalized to nuclear DNA (nucDNA) in BAT from control and CLS-iBKO mice. N=9-13/group. (C) Protein abundance of UCP1, ETS subunits, and CS in whole BAT homogenates. (D) CS activity in whole BAT homogenates. N=4-5/group. (E) Mitochondrial O_2_ consumption (*J*O_2_) in BAT mitochondria from control and CLS-iBKO mice, measured in the presence of 5 mM pyruvate, 0.2 mM malate, 5 mM glutamate, and 5 mM succinate, and 2, 20, and 200 µM ADP. n=5-6/group. (F) ATP production (*J*ATP) in the presence of 5 mM pyruvate, 0.2 mM malate, 5 mM glutamate, 5 mM succinate, and 2, 20, and 200 µM ADP. n=5-6/group. (G) Mitochondrial coupling efficiency (ATP/O ratio). n=5-6/group. (H) UCP1-dependent respiration in BAT mitochondria from control and CLS-iBKO mice. Respiration was stimulated by 5 mM pyruvate and 0.2 mM malate, and UCP1 was subsequently inhibited by 4 mM GDP. n=8-12/group. (I) Protein abundance of UCP1, ETS subunits, and CS in isolated mitochondria. (J) UCP1-dependent respiration stimulated by 0.2 mM malate, 5 mM carnitine, and 20 µM palmitoyl-CoA. n=6/group. (K) UCP1-dependent respiration stimulated by 0.2 mM malate, 5 mM carnitine, and 20 µM palmitoyl-L-carnitine. n=3-5/group. (L) Abundance of phosphorylated (Ser 563 and Ser 660) and total hormone-sensitive lipase (HSL) and adipose triglyceride lipase (ATGL). Data are presented as ± S.E.M. *Denotes of p-value of 0.05 or less.

These findings indicate that, to a substantial extent, CL is dispensable for UCP1 activity. Nevertheless, it is important to acknowledge that CL levels did not reach zero in our CLS-iBKO mice, which leaves open the distinct possibility that the residual CL is sufficient to maintain UCP1 activity in BAT mitochondria from CLS-iBKO mice. We attempted alternative approaches to diminish CL further, but we were unable to achieve this by genetically reducing CLS alone. First, we performed a time-course experiment to see if prolonging terminal experiments would allow CL levels to be more completely depleted. However, new adipocytes began to emerge between 6- and 8-weeks post-tamoxifen injection, preventing mitochondrial CL from reaching zero. We also attempted to reduce mitochondrial CL levels to zero in vitro by culturing primary brown adipocytes. However, deleting CLS in pre-adipocytes prevented brown adipocyte differentiation, and deleting CLS post-differentiation did not lower mitochondrial CL beyond the level that was achievable in vivo. Thus, we were unable to conclude that CL is completely dispensable for UCP1 activity. Nevertheless, it is important to point out that CLS-iBKO mice exhibited cold intolerance and reduced VO_2_ induced by β3-agonist (Figure 2H-J) despite incomplete removal of CL. Therefore, it can be concluded that defective thermogenesis in CLS-iBKO mice is not exclusively due to the direct effect of CL on UCP1. In turn, our findings suggest that CLS influences thermogenesis independent of its action on UCP1, including the possibility that CLS may have an alternate enzyme activity in addition to the synthesis of CL.

How does CLS deletion impair thermogenesis independent of modulating UCP1 activity? Control and CLS-iBKO mice did not differ in UCP1, complex I-V, or citrate synthase content per unit of mitochondria (Figure 3I). Combined with unchanged total cellular UCP1 content (Figure 3C), these data show that reduced thermogenesis in CLS-iBKO mice is not due to CLS regulating UCP1 protein level. We then examined the possibility that CLS affects the ability of fatty acids to activate UCP1.

However, respiration driven by palmitoyl-CoA was not different between the groups (Figure 3J). Based on the possibility that low CL may interfere with the transport of fatty acids across OMM and IMM (Pande, 1975; Violante et al., 2013), we also tested respiration induced by palmitoyl-L-carnitine. However, similar to respiration driven by palmitoyl-CoA, CLS deletion had no effect on respiration driven by palmitoyl-L-carnitine (Figure 3K). These data suggest that CL deletion does not interfere with the ability of fatty acids to activate UCP1, nor does it affect the ability of the carnitine system to transport fatty acids across mitochondrial membranes. Lastly, we examined the possibility that CL influences the availability of fatty acids necessary to activate UCP1. BAT thermogenesis is activated in vivo by noradrenaline binding to the β3 G-protein coupled receptor (GPCR) on brown adipocytes. The signaling cascade activates hormone sensitive lipase (HSL) and adipose triglyceride lipase (ATGL), both of which are required to cleave triacylglycerol into individual free fatty acids (FA) (Cannavo and Koch, 2017; Schena and Caplan, 2019). Deletion of CLS did not lower the protein abundance of either HSL or ATGL, nor affect two activating phosphorylation sites on HSL (Figure 3L) (Anthonsen et al., 1998; Holm, 2003; Holm et al., 1997). Thus, the role of CLS in regulating BAT thermogenesis is independent of fatty acid transport and signaling as well as fatty acid-mediated UCP1 activation.

### Mitochondrial PE is essential for brown adipose thermogenesis

Mitochondria have the intrinsic ability to modulate the abundance of PE by an IMM-resident enzyme phosphatidylserine decarboxylase (PSD) (Figure 4A) (Heden et al., 2019; van der Veen et al., 2017; Vance and Tasseva, 2013). A mitochondrial importer for PE is not known to exist, and stable isotope studies suggest that PE is not imported into mitochondria (Shiao et al., 1995). Thus, mitochondrial PE is likely almost exclusively generated by PSD. Because mitochondrial PE was the most affected class of lipids that were energy-responsive in BAT, we developed a system whereby we could manipulate the levels of mitochondrial PE and test its role in BAT thermogenesis. To this end, we generated mice with tamoxifen-inducible UCP1-Cre driven knockout of PSD (PSD-iBKO) to block mitochondrial PE synthesis (Figure 4B&C).

**Figure 4:**
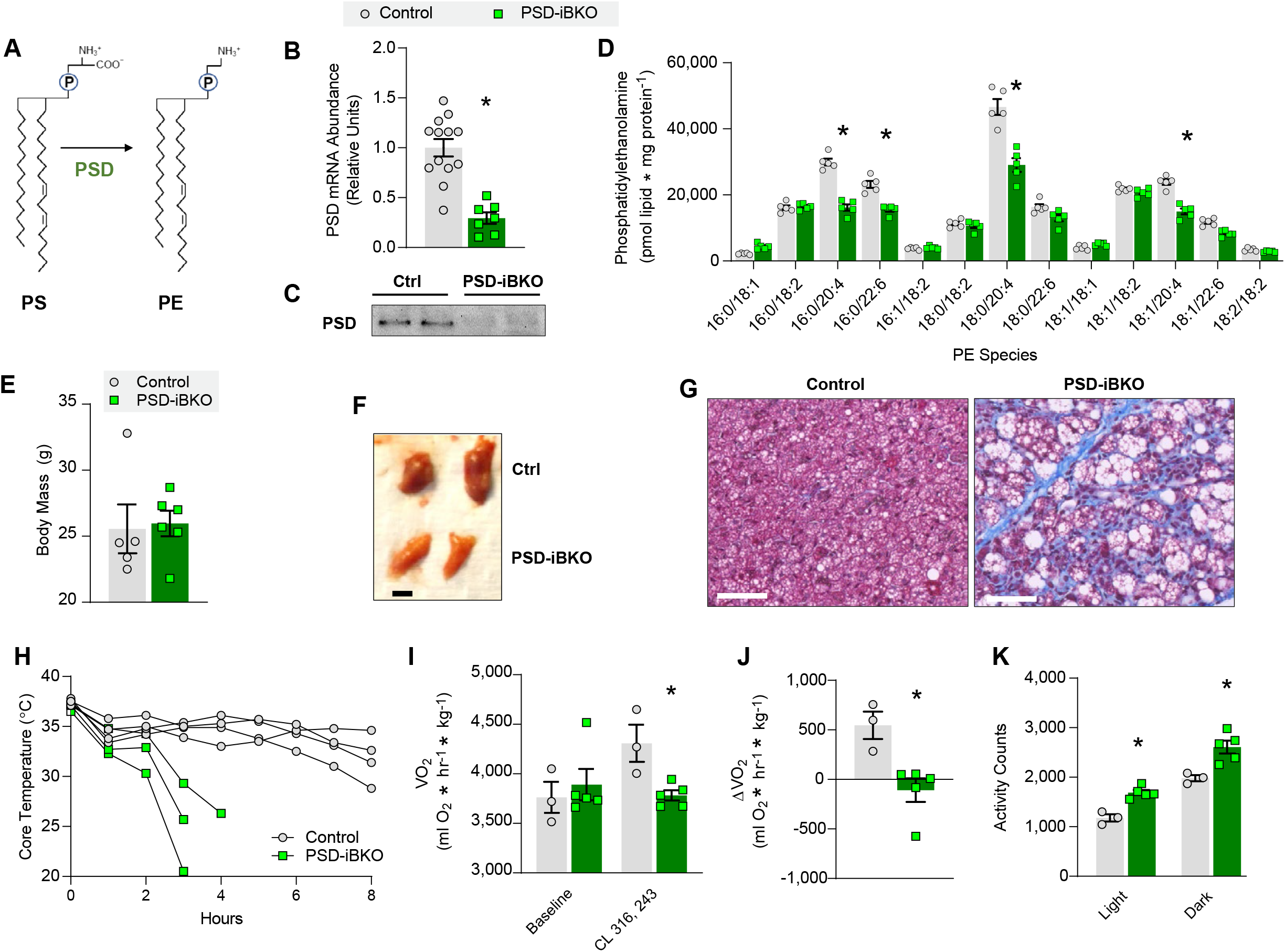
Loss of mitochondrial PE impairs brown adipose thermogenesis. (A) A schematic for mitochondrial PE biosynthesis. (B) PSD mRNA abundance in BAT from control and PSD-iBKO mice. n=10-12/group. (C) Representative images of PSD western blot in BAT from control and PSD-iBKO mice. (D) Mitochondrial PE species abundance in BAT 2 weeks post-tamoxifen injection. n=5/group. (E) Body mass. n=5-6/group. (F) Representative intrascapular BAT depots from control and PSD-iBKO mice. (G) Masson’s trichrome blue staining of BAT. Scale bar = 50 µM. (H) Core body temperature of mice subjected to an acute cold-tolerance test at 4° C. n=3-4/group. (I) Whole-body oxygen consumption in mice before and after injection of CL-316,243. n=3-5/group. (J) Changes in whole-body oxygen consumption following administration of CL-316,243. n=3-5/group. (K) Spontaneous movement in metabolic cages. n=3-5/group. Data are presented as ± S.E.M. *Denotes of p-value of 0.05 or less.

A recent report suggests that constitutive UCP1-Cre mice have leaky Cre expression in other key metabolic tissues such as the hypothalamus and kidney resulting in nonspecific genetic recombination (Claflin et al., 2022). We examined whether the tamoxifen-inducible UCP1-Cre (UCP1 Cre-ERT2) we used resulted in PSD knockout in other tissues. Although we did detect a small level of Cre expression in the hypothalamus and some other non-adipose tissues, PSD expression was not significantly altered in any tissue but BAT (Figure S5A&B). These results suggest that any observed mitochondrial or thermogenic phenotypes in PSD-iBKO mice are unlikely to result from changes in PSD expression in cell types other than brown or beige adipocytes.

Unlike CLS-iBKO mice, which were studied 4-weels post-tamoxifen injection, the level of mitochondrial PE was reduced 2-weeks post-tamoxifen injection in PSD-iBKO mice (Figure 4D), demonstrating a more rapid turnover of PE than CL. Extending the terminal experiments to 4-week post-tamoxifen injection did not further lower mitochondrial PE level (Figure S6A), likely due to adipocyte turnover as well as compensatory PE synthesis by mitochondrial lyso-PE acylation (Lands cycle) (Figure S6B). There were also changes in the levels of mitochondrial PS (the substrate of PSD), which further demonstrated loss of PSD activity (Figure S6C). Thus, we performed all our subsequent experiments on PSD-iBKO mice 2-4 weeks post-tamoxifen injection.

Reduction of PSD driven by UCP1 Cre-ERT2 did not appear to have an overt phenotype in an unstressed condition; for instance, control and PSD-iBKO mice did not differ in body mass (Figure 4E). At the tissue level, however, BAT from PSD-iBKO mice were smaller and paler compared to control mice (Figure 4F, S6D). This was in contrast to BAT from CLS-iBKO mice that were larger compared to their controls. Histological analyses revealed larger lipid droplets as well as fibrosis in BAT from PSD-iBKO mice (Figure 4G). Intracellular lipid accumulation may suggest reduced thermogenic capacity (Alcalá et al., 2017; Turchi et al., 2020). Indeed, PSD-iBKO mice demonstrated substantial reduction in cold tolerance compared to controls (Figure 4H). Furthermore, PSD-iBKO mice were not responsive to CL-316,243-induced increase whole-body oxygen consumption (Figure 4I&J), suggesting impaired BAT thermogenesis. Potentially as a mechanism to compensate for the lack of brown adipose thermogenesis, PSD-iBKO mice were more active than control mice during light and dark cycles (Figure 4K). This coincided with trends for elevated whole-body oxygen consumption and RER, with PSD deletion (Figure S6D&E).

### Mitochondrial PE is required for proton flux through UCP1

To investigate the mechanistic role of mitochondrial PE in thermogenesis, we examined BAT mitochondria from control and PSD-iBKO mice. Electron micrograph revealed robust disorganization of IMM, particularly with decreased density of cristae (Figure 5A). Quantification of mtDNA/nucDNA or mitochondrial enzymes by western blotting revealed that PSD deletion substantially lowered mitochondrial density per cell (Figure 5B&C). In addition to reduced mitochondrial density, PSD deletion also lowered the oxidative capacity per unit of mitochondria. High-resolution respirometry and fluorometry of isolated mitochondria from BAT revealed reduced ADP-dependent respiration in PSD-iBKO mice compared to controls (Figure 5D). Strikingly, the reduction was not due to changes in the rate of ATP synthesis (Figure 5E) or electron leak (Figure S6F). Instead, increased ATP/O ratio (Figure 5F) coincided with a robust reduction in UCP1-dependent respiration (Figure 5G). Importantly, the reduction in UCP1-dependent respiration occurred in the absence of changes in UCP1 protein abundance per unit of mitochondria (Figure 5H). Consistent with unaltered ATP synthesis, abundances of Complex I-V in mitochondria were not different between control and PSD-iBKO mice. These findings suggest that mitochondrial PE is essential for optimal UCP1 function.

**Figure 5:**
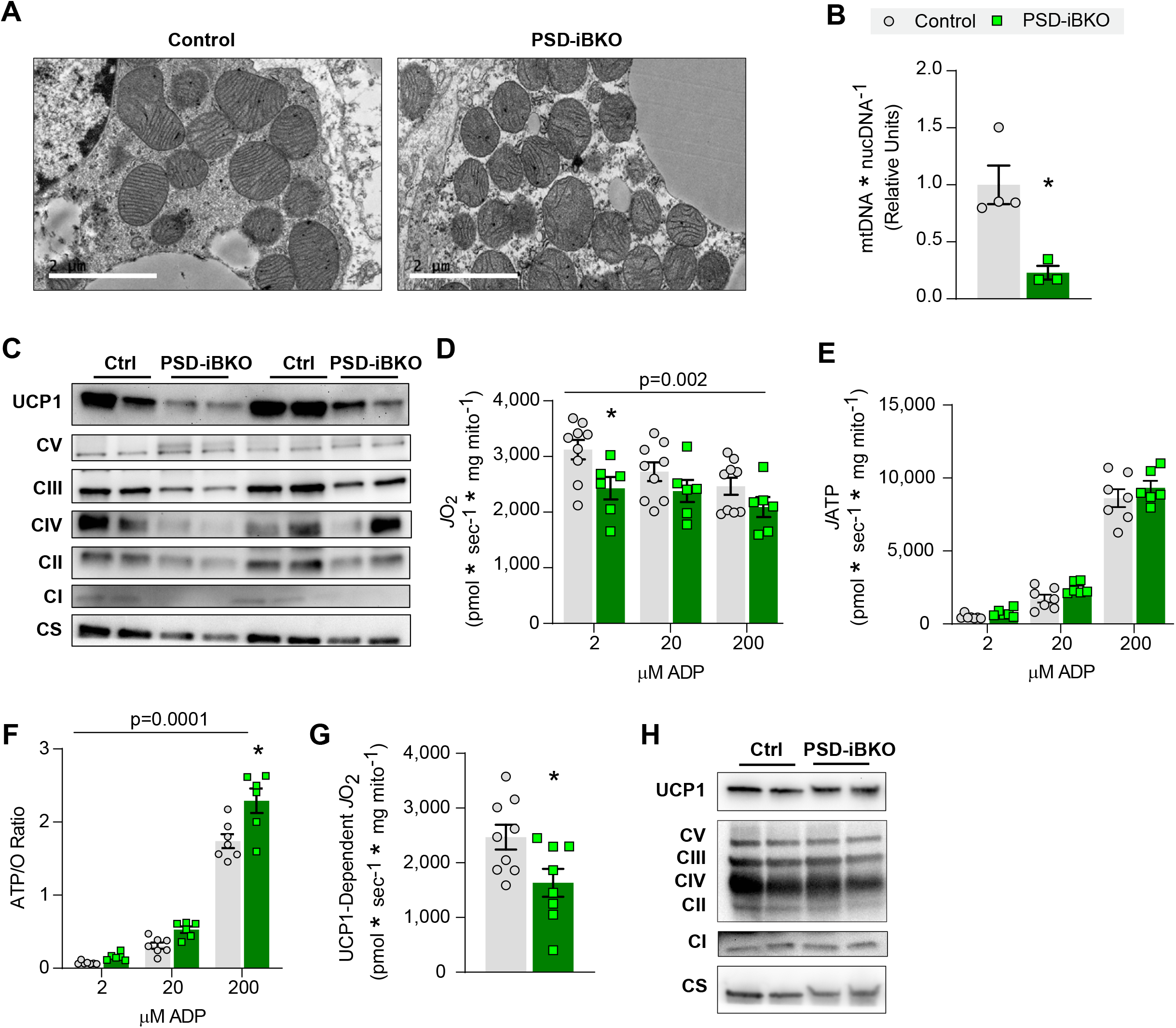
Mitochondrial PE is essential for UCP1-dependent respiration. (A) Transmission electron microscopy images of BAT mitochondria from control and PSD-iBKO mice. Scale bar=2 µM. (B) Mitochondrial DNA levels (mtDNA) normalized to nuclear DNA (nucDNA) in BAT from control and PSD-iBKO mice. n=3-4/group. (C) Protein abundance of UCP1, ETS subunits, and CS in whole BAT homogenates. (D) Mitochondrial O_2_ consumption in BAT mitochondria from control and PSD-iBKO mice, measured in the presence of 5 mM pyruvate, 0.2 mM malate, 5 mM glutamate, and 5 mM succinate, and 2, 20, and 200 µM ADP. n=8-9/group. (E) ATP production in the presence of 5 mM pyruvate, 0.2 mM malate, 5 mM glutamate, 5 mM succinate, and 2, 20, and 200 µM ADP. n=8-9/group. (F) Mitochondrial coupling efficiency (ATP/O ratio). n=8-9/group. (G) UCP1-dependent respiration in BAT mitochondria from control and PSD-iBKO mice. Respiration was stimulated by 5 mM pyruvate and 0.2 mM malate, and UCP1 was subsequently inhibited by 4 mM GDP. n=8-9/group. (H) Protein abundance of UCP1, ETS subunits, and CS in isolated mitochondria. Data are presented as ± S.E.M. *Denotes of p-value of 0.05 or less.

To more directly assess the effect of PE on UCP1 activity, we quantified UCP1-dependent proton current in mitoplasts prepared from BAT mitochondria in control and PSD-iBKO mice (Figure 6A) (Balderas et al., 2022; Fedorenko et al., 2012). IMM portion of mitoplasts were patch-clamped to perform electrophysiologic measurements of proton current through UCP1 (Figure 6B). The UCP1 contribution to the proton current density is the portion of the total baseline current that is inhibited by the subsequent addition of 1 mM ATP (Figure 6C&D). Strikingly, total proton current was substantially and consistently diminished in BAT mitoplasts from PSD-iBKO mice compared to control mice, whereas the difference was abolished after addition of ATP (Figure 6E). Quantifying UCP1 proton current density as the difference between baseline and ATP addition revealed 60% lower levels in mitoplasts from PSD-iBKO mice compared to control mice (Figure 6F). Thus, while UCP1 levels within mitochondria were unchanged, its activity was clearly reduced, suggesting that mitochondrial PE can facilitate UCP1 activity.

**Figure 6:**
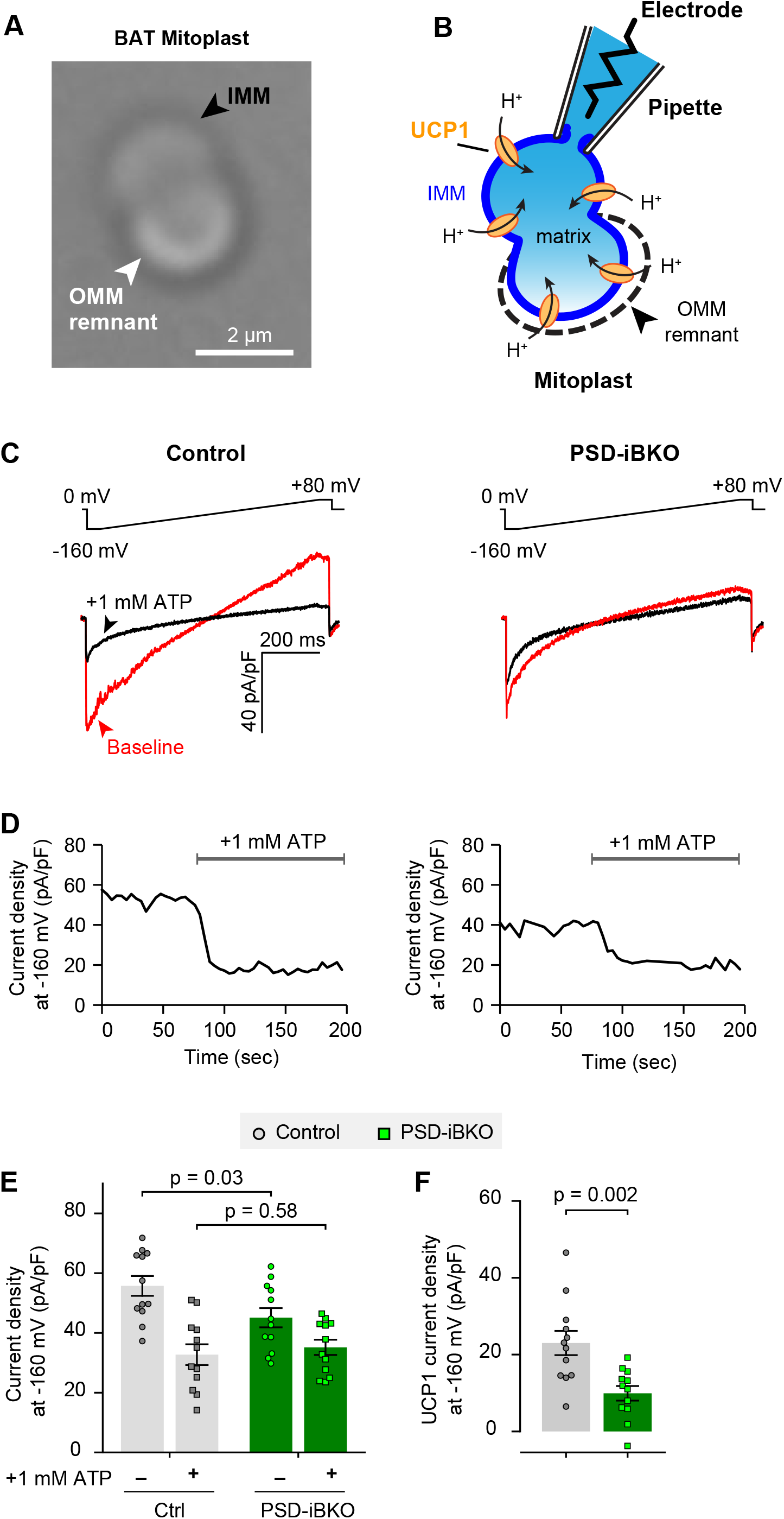
Mitochondrial PE is essential for proton current through UCP1. (A) Differential interference contrast image of BAT mitoplasts reveals typical bi-lobed appearance. IMM: black arrow. OMM remnant: white arrow. Scale bar= 2 µM. (B) Schematic illustrating electrophysiological recording setup for proton (H^+^) current. (C) Top, voltage ramp protocol. Bottom, exemplar traces showing baseline proton currents (red) and after addition of 1 mM ATP (black). Measurements are taken at - 160 mV (arrowheads). (D) Exemplar time course of proton current inhibition with ATP. (E) Summary of proton current densities (n=12 mitoplasts/group). (F) Quantification of UCP1 current density, taken as the difference for each mitoplast between baseline and after ATP addition, from (E) (n=12 mitoplasts/group). Data are presented as ± S.E.M.

## DISCUSSION

OXPHOS is the process whereby energy derived from substrates is transduced by a series of reactions that occur in and across the IMM to ultimately yield ATP synthesis. UCP1, the chief enzyme of brown adipose thermogenesis, also resides in IMM where it disrupts the proton gradient to uncouple ATP synthesis from ETS. Here, we provide evidence that mitochondrial PE is an energy-responsive IMM metabolite that alters UCP1 activity to regulate thermogenesis. At the organism level, mice with reduced mitochondrial PE were cold intolerant and insensitive to β3-agonist induced increase in whole-body oxygen consumption. Deep phenotyping of BAT mitochondria revealed that low PE robustly reduces UCP1-dependent respiration and UCP1 proton current. Together, these findings suggest that mitochondrial PE responds to changes in BAT thermogenic demand to potentiate UCP1 activity.

Previously, we demonstrated that adipose CL is essential for BAT thermogenesis and systemic energy homeostasis (Sustarsic et al., 2018). CL has been implicated in mitochondrial function in various tissues including liver, skeletal muscle, and BAT (Lynes et al., 2018; Ostojic et al., 2013; Paradies et al., 2014; 2019). As a lipid that is almost exclusively localized in IMM, it would be reasonable to suspect that CL might directly affect UCP1 function to uncouple OXPHOS. Indeed, some studies suggest that CL stabilizes UCP1 (Lee et al., 2015), while other studies suggest CL attenuates inhibition of UCP1 by nucleotide binding (Klingenberg, 2009). However, we found that although CLS deletion indeed made mice cold intolerant, it had no effect on UCP1-dependent respiration. Nonetheless, it is important to acknowledge that our strategy with CLS knockout did not achieve a complete deprivation of CL in the IMM milieu. Mitochondrial CL level also did not decrease with thermoneutrality or UCP1 knockout, suggesting that CL level does not universally respond to changes in BAT thermogenic burden. Indeed, our unpublished data in non-adipose tissues suggest that CL acutely responds to changes in diet, and cold exposure is known to increase food intake (Jia et al., 2016; Toloza et al., 1991).

What is the mechanism by which CL regulates thermogenesis independent of UCP1? In our studies, we ruled out an effect of CL on BAT mitochondrial density, cristae morphology, or UCP1 protein abundance. We also ruled out the possibility that CL alters the sensitivity of UCP1 to be activated by fatty acids, or changes in GPCR signaling that induces fatty acid mobilization. Our previous study suggests that CLS has a robust effect on modulating transcription (Sustarsic et al., 2018), and we continue to subscribe to this idea that CL or CLS may have an alternate role to regulate metabolism independent of mitochondria.

Unlike CL, mitochondrial PE responded to cold, thermoneutrality, and UCP1 knockout, making this lipid a candidate for an energy-responsive UCP1 rheostat. Indeed, decreased mitochondrial PE directly reduced UCP1 proton current, without altering electron leak or ATP synthesis. Importantly, the reduction in UCP1 activity occurred in the absence of changes in UCP1 protein content per unit of mitochondria. This suggests that PE directly activates UCP1 function. How does mitochondrial PE regulate UCP1? Unlike CL, PE has not been implicated to bind to UCP1 (Lee et al., 2015). However, a study with UCP1-reconstituted liposomes suggests that PE enhances the UCP1 proton translocation (Jovanovic et al., 2015). The proposed mechanism was that PE adducts alter the membrane boundary potential of the IMM bilayer to facilitate the protonophoric activity of UCP1. We note that reduced levels of PE coincided with robustly deformed cristae, an observation consistent with their model on the effect that PE has on membrane curvature. Regardless of the mechanism, compromised UCP1 function with low mitochondrial PE was sufficient to impair thermogenesis induced by cold or β3-agonist administration. Taken together with our data that cold or thermoneutrality regulates the concentration of PE in mitochondria, we postulate that PE-dependent regulation of UCP1 function is a physiologically-relevant mechanism for thermogenesis.

We also attempted to increase mitochondrial PE content to see if it would be sufficient to increase UCP1 function. In summary, both our in vivo and in vitro approaches failed to sufficiently increase mitochondrial PE in brown adipocytes. For in vivo experiments, we performed PSD overexpression by examining mice with conditional PSD overexpression using UCP1-Cre (PSD-BKI, Figure S7A-M). We previously used this strategy to successfully increase PSD expression and mitochondrial PE in skeletal muscle (Heden et al., 2019). While PSD transcript was successfully elevated in BAT from PSD-BKI mice compared to control mice, this intervention only increased mitochondrial PE by ∼10%, likely due to endogenous PSD activity that is already high in BAT. Further, we did not observe any differences in β3-agonist induced oxygen consumption or UCP1-dependent respiration. Similarly, in vitro ethanolamine supplementation was not effective in increasing mitochondrial PE content, and doses of lyso-PE supplementation that were sufficient to increase mitochondrial PE also induced cell death. Lastly, we performed experiments using small unilamellar vesicles (SUVs) in an attempt to directly deliver PE to mitochondria. However, PE alone does not effectively form SUVs, and fusion of PE/PC SUVs at various ratios all substantially diluted mitochondrial proteins and lowered total mitochondrial respiration. Thus, at this point in time, we are unable to provide evidence for gain-of-function of PSD or mitochondrial PE to increase UCP1 function.

How do changes in temperature regulate PE and other mitochondrial lipids? Our data suggest that mitochondrial PE is at least partly regulated by changes in PSD transcription. There is very little known regarding the transcriptional control of enzymes for mitochondrial lipid biosynthesis and transport. In skeletal muscle, we previously showed that exercise or sedentariness influences PSD to regulate mitochondrial PE level (Heden et al., 2019). Thus, we suspect that the PSD-mitochondrial PE axis bidirectionally responds to changes in energy demand to optimize mitochondrial bioenergetics in multiple cell-types. Conversely, we suspect that the CLS-mitochondrial CL axis responds to changes in energy supply. Dietary interventions such as high-fat diet feeding and calorie restriction are known to influence cellular CL levels in multiple tissues (Feillet-Coudray et al., 2014; He and Han, 2014; Luevano-Martinez et al., 2017; Sullivan et al., 2017; Zhang et al., 2022). We have preliminary evidence in non-adipose tissues that mitochondria CL specifically responds to diet interventions. In the current study, cold intervention increased mitochondrial CL content, which is known to also increase food intake in mice (Jia et al., 2016; Smith and Romsos, 1984; Toloza et al., 1991). These lines of evidence provide important insights into a more global understanding of how mitochondrial lipids such as PE and CL respond to energy demand or supply to influence mitochondrial energetics (Figure 7).

**Figure 7:**
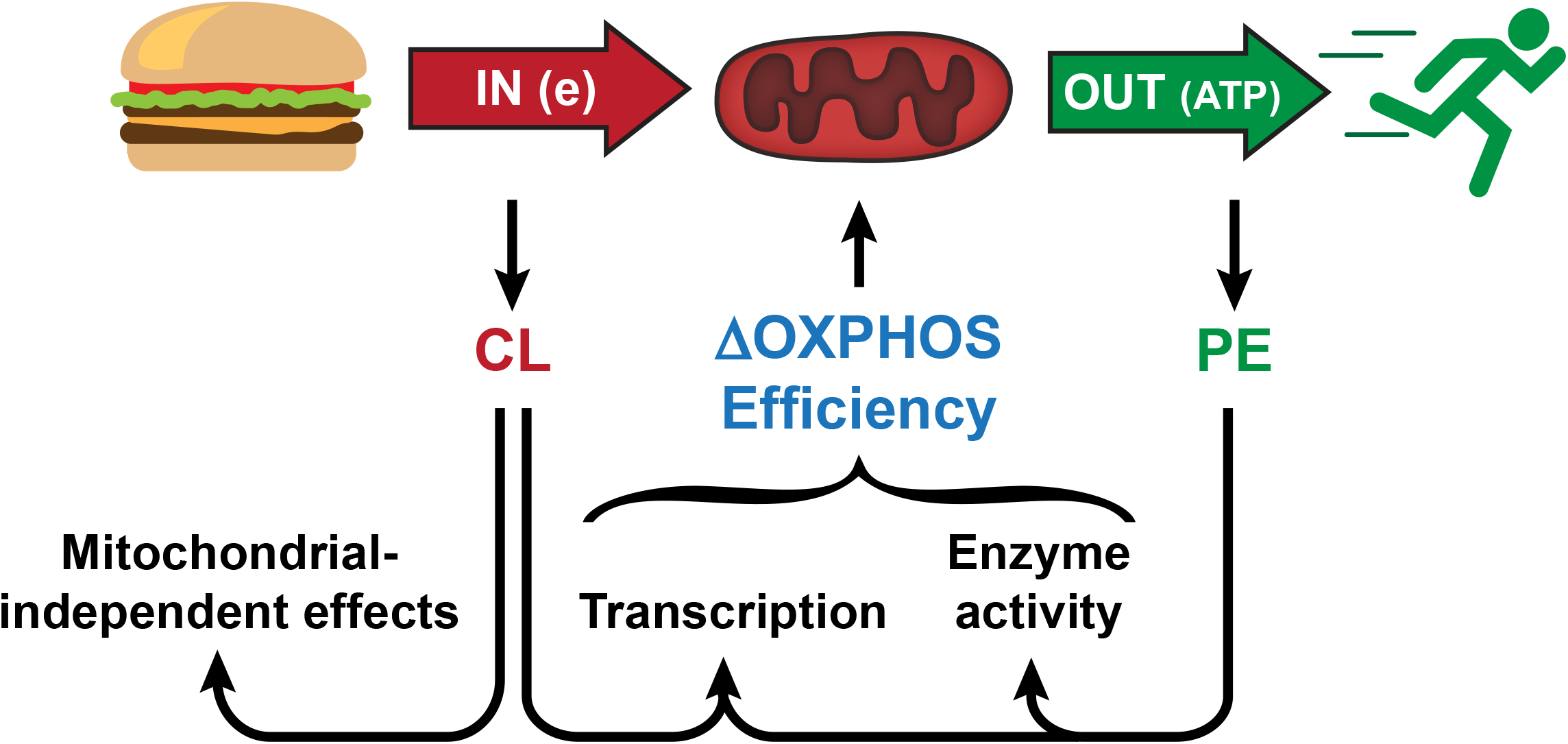
Mitochondrial phospholipids as OXPHOS rheostat that respond to energy demand or supply. We propose that mitochondria respond to changes in energy demand or supply by altering the levels of mitochondrial PE and CL, respectively. We postulate that these lipids in turn influence mitochondrial content and OXPHOS efficiency, through distinct but potentially universal mechanisms in mammalian cells.

Harnessing UCP1-dependent thermogenesis remains an attractive strategy for treating obesity and/or hyperglycemia. In this study, we identified an important role that mitochondrial PE plays in UCP1-dependent thermogenesis in BAT. CL did not appear to directly regulate UCP1 function, although it is clear that the CLS-CL axis plays a role in regulating thermogenesis independent of mitochondrial UCP1 activity. Combined with our previous findings, we propose that mitochondrial PE is a universal cellular rheostat that modulates mitochondrial efficiency in response to changes in energy demand.

## ACKNOWLEDGEMENTS

This work was supported by funding from NIH grants DK107397, DK127979, GM144613, AG074535 (K.F.), DK130555 (A.D.P), DK091317 (M.J.L.), DK103930 (C.J.V.), DK115867, DK132239 (I.J.L.), GM131854, CA228346 (J.R.), HL141353, HL165797 (D.C.), Howard Hughes Medical Institute (J.R.), Jane Coffin Childs Memorial Fund (J.T.M.), Nora Eccles Treadwell Foundation (D.C.), Department of Defense W81XWH-19-1-0213 (to K.F-W), and American Heart Association grant 19PRE34380991 (J.M.J.). University of Utah Metabolomics Core Facility is supported by S10 OD016232, S10 OD021505, and U54 DK110858. We thank Diana Lim from University of Utah Molecular Medicine for assistance with figures.

## AUTHOR CONTRIBUTIONS

J.M.J., A.D.P., Z.G-H., and K.F. conceived the project and designed the experiments. J.M.J. and A.D.P. conducted the majority of experiments for this manuscript. E.G.S., V.P., C.J.V., and Z.G-H. assisted with generation of mouse models. J.A.M. and J.E.C. conducted lipidomic mass spectrometry. E.B., M.J.L., A.J-R., K.H.F-W., and D.C. assisted with mitochondrial energetics experiments. J.T.M., J.R., A.S., I.J.L., K.H.F-W., and D.C. provided assistance with experimental design. A.D.P. and K.F. wrote the manuscript. This manuscript was reviewed by all authors, revised, and given approval by all authors for publication.

## FIGURE LEGENDS

**Supplemental Figure 1:**
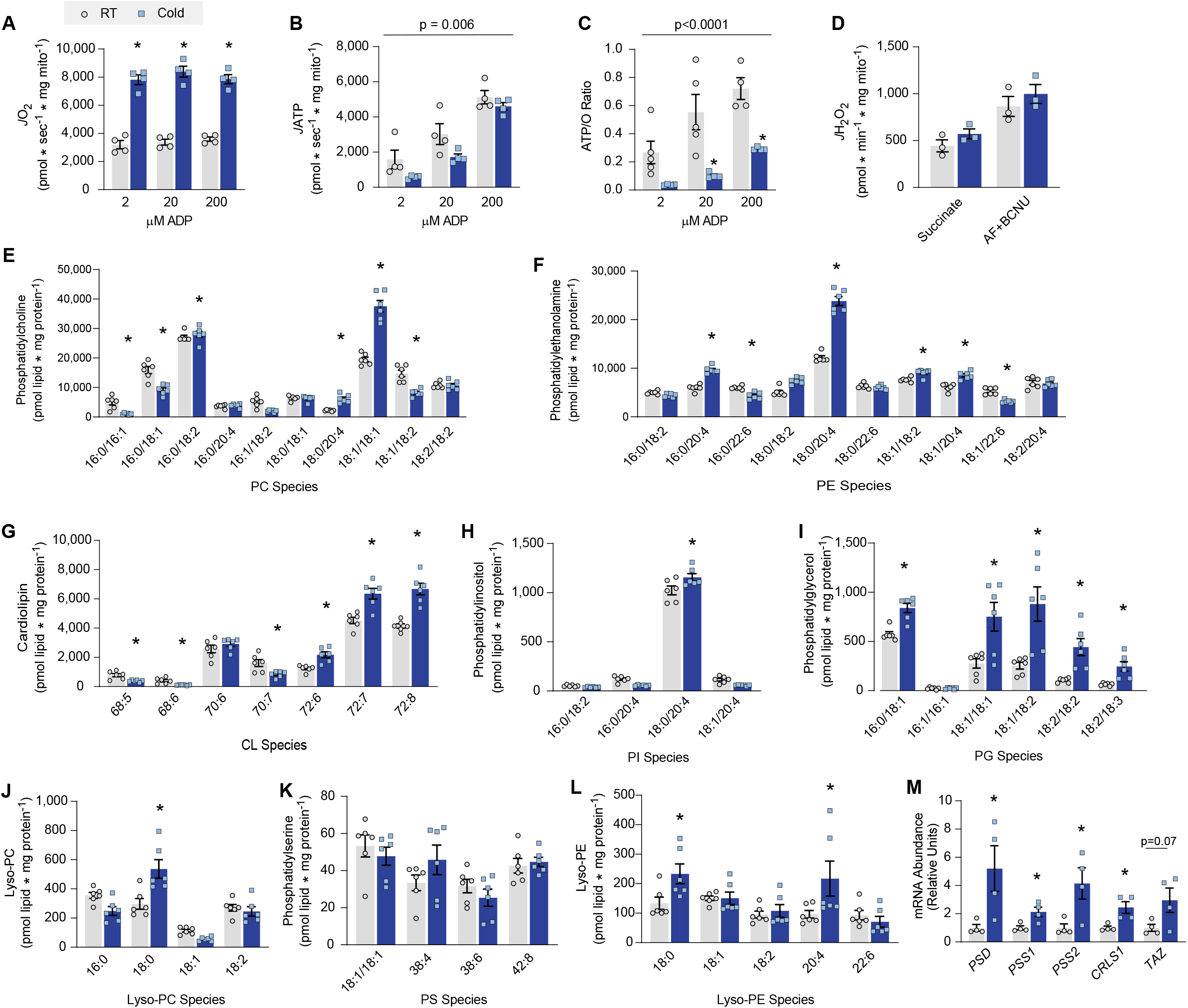
Cold-induced adaptations in mitochondrial bioenergetics and phospholipids in BAT. (A) Mitochondrial O_2_ consumption in BAT from C57BL/6J mice housed at RT or 6.5 °C for 7 days, measured in the presence of 5 mM pyruvate, 0.2 mM malate, 5 mM glutamate, 5 mM succinate, and 2, 20, and 200 µM ADP from. n=4/group. (B) ATP production in the presence of 5 mM pyruvate, 0.2 mM malate, 5 mM glutamate, 5 mM succinate, and 2, 20, and 200 µM ADP. n=4/group. (C) Mitochondrial coupling efficiency (ATP/O ratio) of BAT mitochondria in C57BL/6J mice housed at RT or 6.5 °C for 7 days. n=4/group. (D) H_2_O_2_ production in BAT mitochondria from C57BL/6J mice housed at RT and 6.5 °C, measured in the presence of 10 mM succinate and antioxidant inhibitors auranofin (AF) and carmustine (BCNU). n=4/group. (E-L) Mass spectrometric analyses of mitochondrial lipids in BAT from C57BL/6J mice housed at RT and 6.5 °C. PC (E), PE (F), CL (G), PI (H), PG (I), lyso-PC (J), PS (K), and lyso-PE (L). n=6/group. (M) mRNA abundance of genes of mitochondrial PE and CL biosynthesis. n=4/group. Data are presented as ± S.E.M. *Denotes of p-value of 0.05 or less.

**Supplemental Figure 2:**
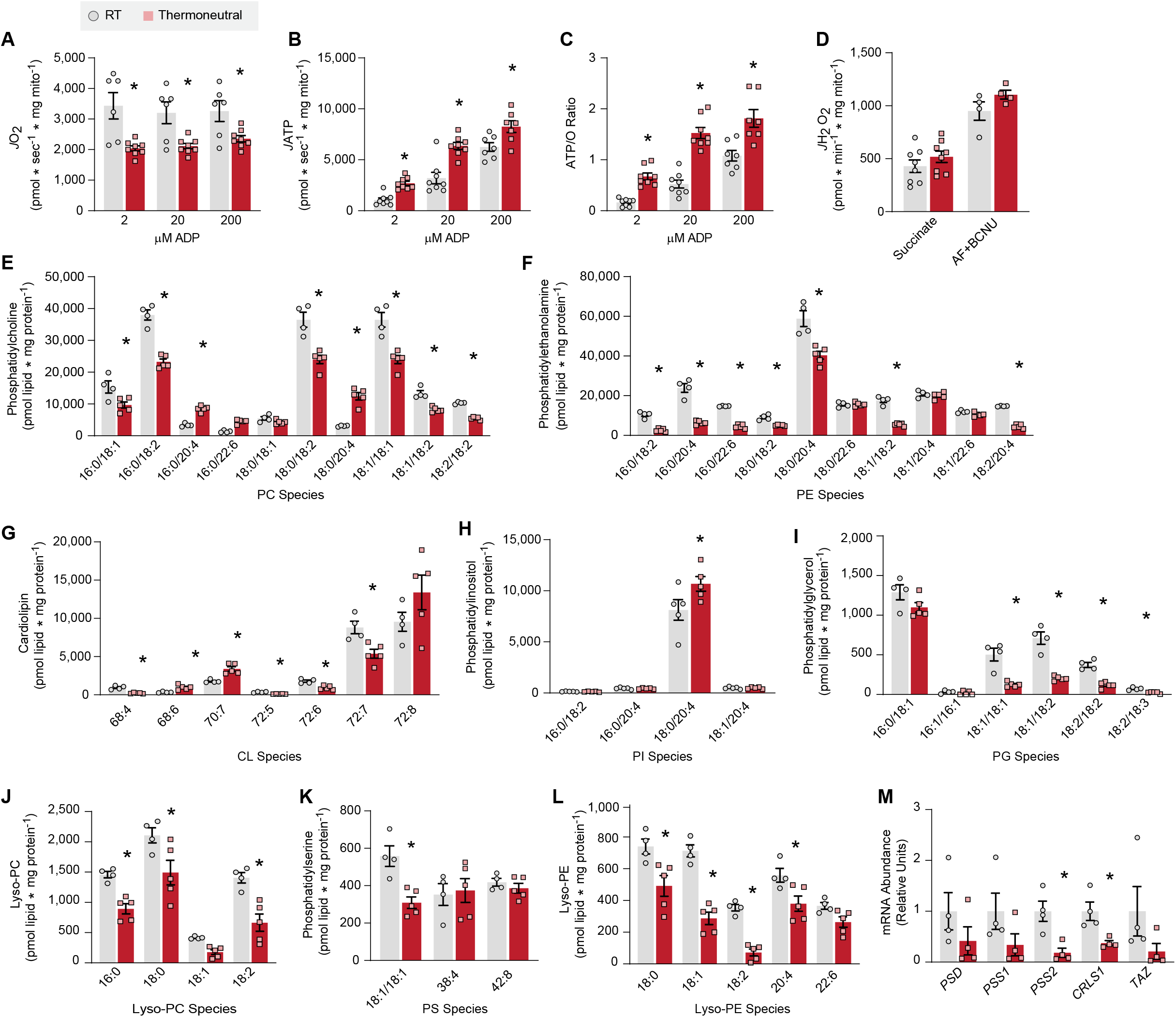
Thermoneutrality-induced adaptations in mitochondrial bioenergetics and phospholipids in BAT. (A) Mitochondrial O_2_ consumption in BAT from C57BL/6J mice housed at RT or 30 °C for 30 days, measured in the presence of 5 mM pyruvate, 0.2 mM malate, 5 mM glutamate, 5 mM succinate, and 2, 20, and 200 µM ADP from. n=8/group. (B) ATP production in the presence of 5 mM pyruvate, 0.2 mM malate, 5 mM glutamate, 5 mM succinate, and 2, 20, and 200 µM ADP. n=8/group. (C) Mitochondrial coupling efficiency (ATP/O ratio) of BAT mitochondria. n=8/group. (D) H_2_O_2_ production in the presence of 10 mM succinate and antioxidant inhibitors auranofin (AF) and carmustine (BCNU). n=4-8/group. (E-L) Mass spectrometric analyses of mitochondrial lipids in BAT from C57BL/6J mice housed at RT and 30 °C. PC (E), PE (F), CL (G), PI (H), PG (I), lyso-PC (J), PS (K), and lyso-PE (L). n=4-5/group. (M) mRNA abundance of genes of mitochondrial PE and CL biosynthesis. n=4/group. Data are presented as ± S.E.M. *Denotes of p-value of 0.05 or less.

**Supplemental Figure 3:**
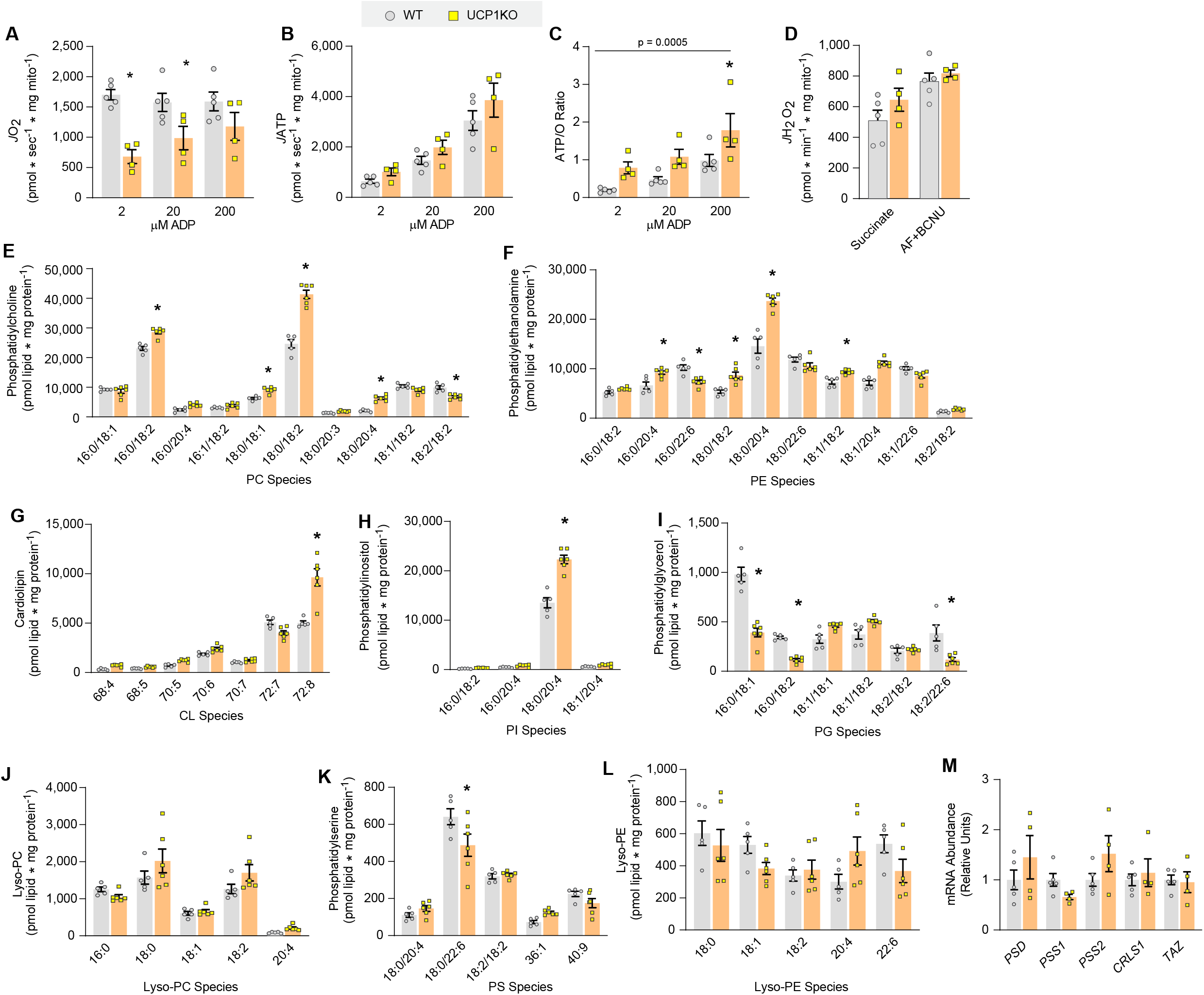
Effects of UCP1 deletion on mitochondrial bioenergetics and phospholipids in BAT. (A) Mitochondrial O_2_ consumption in BAT from WT and UCP1KO mice, measured in the presence of 5 mM pyruvate, 0.2 mM malate, 5 mM glutamate, 5 mM succinate, and 2, 20, and 200 µM ADP. n=4-5/group. (B) ATP production in the presence of 5 mM pyruvate, 0.2 mM malate, 5 mM glutamate, 5 mM succinate, and 2, 20, and 200 µM ADP. n=4-5/group. (C) Mitochondrial coupling efficiency (ATP/O ratio) of BAT mitochondria. n=4-5/group. (D) H_2_O_2_ production in the presence of 10 mM succinate and antioxidant inhibitors auranofin (AF) and carmustine (BCNU). n=4-5/group. (E-L) Mass spectrometric analyses of mitochondrial lipids in BAT from WT and UCP1KO mice. PC (E), PE (F), CL (G), PI (H), PG (I), lyso-PC (J), PS (K), and lyso-PE (L). n=5-6/group. (M) mRNA abundance of genes of mitochondrial PE and CL biosynthesis. n=4/group. Data are presented as ± S.E.M. *Denotes of p-value of 0.05 or less.

**Supplemental Figure 4:**
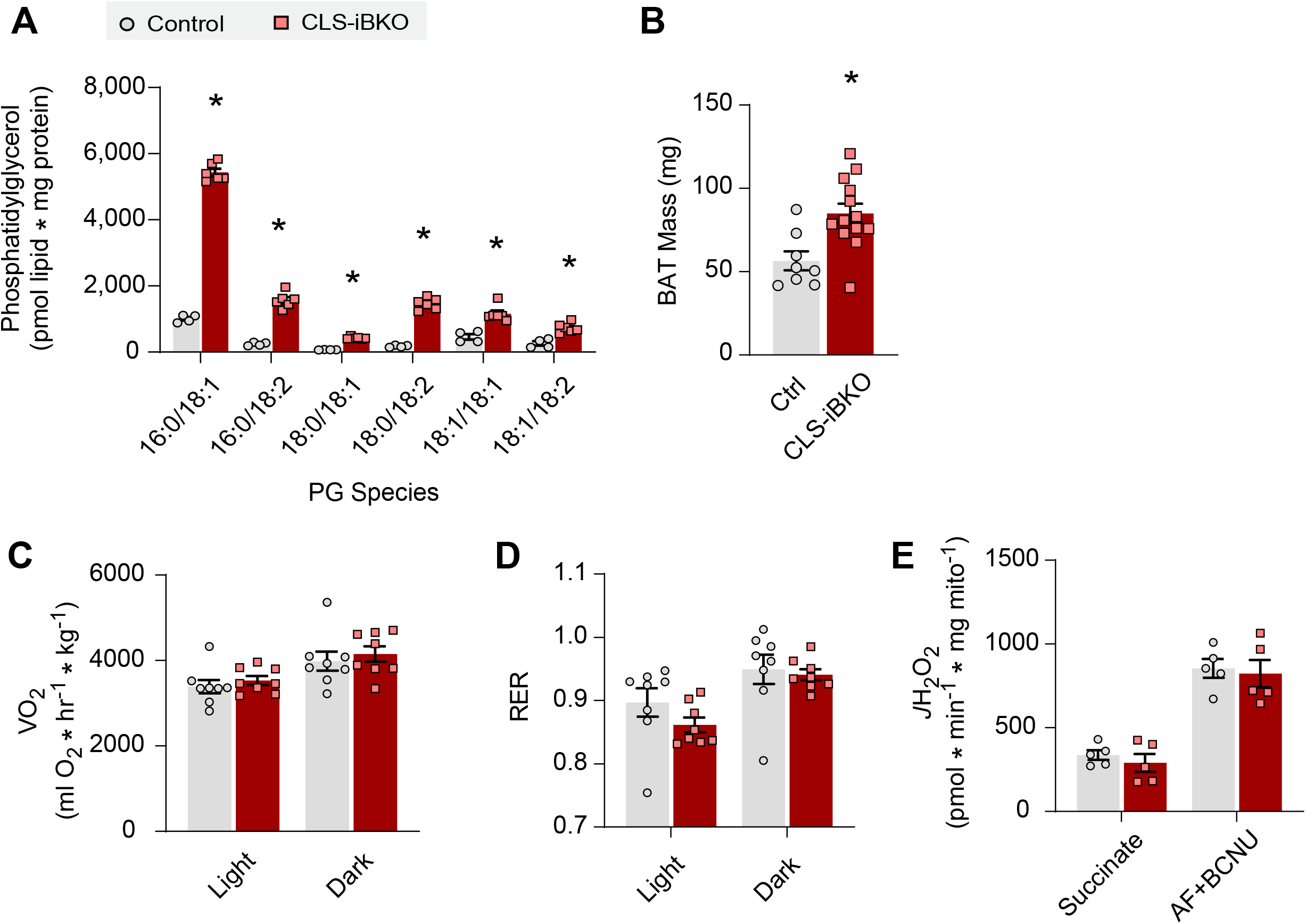
Additional data on CLS-iBKO mice. (A) Mitochondrial PG, a precursor to mitochondrial CL. n=4-6/group. (B) BAT mass. n=8-13/group. (C) Whole-body oxygen consumption in metabolic cage. n=8/group. (D) Respiratory exchange ratio (RER) in metabolic cage. n=8/group. (E) Mitochondrial H_2_O_2_ production in the presence of 10 mM succinate and antioxidant inhibitors auranofin (AF) and carmustine (BCNU). n=5/group. Data are presented as ± S.E.M. *Denotes of p-value of 0.05 or less.

**Supplemental Figure 5:**
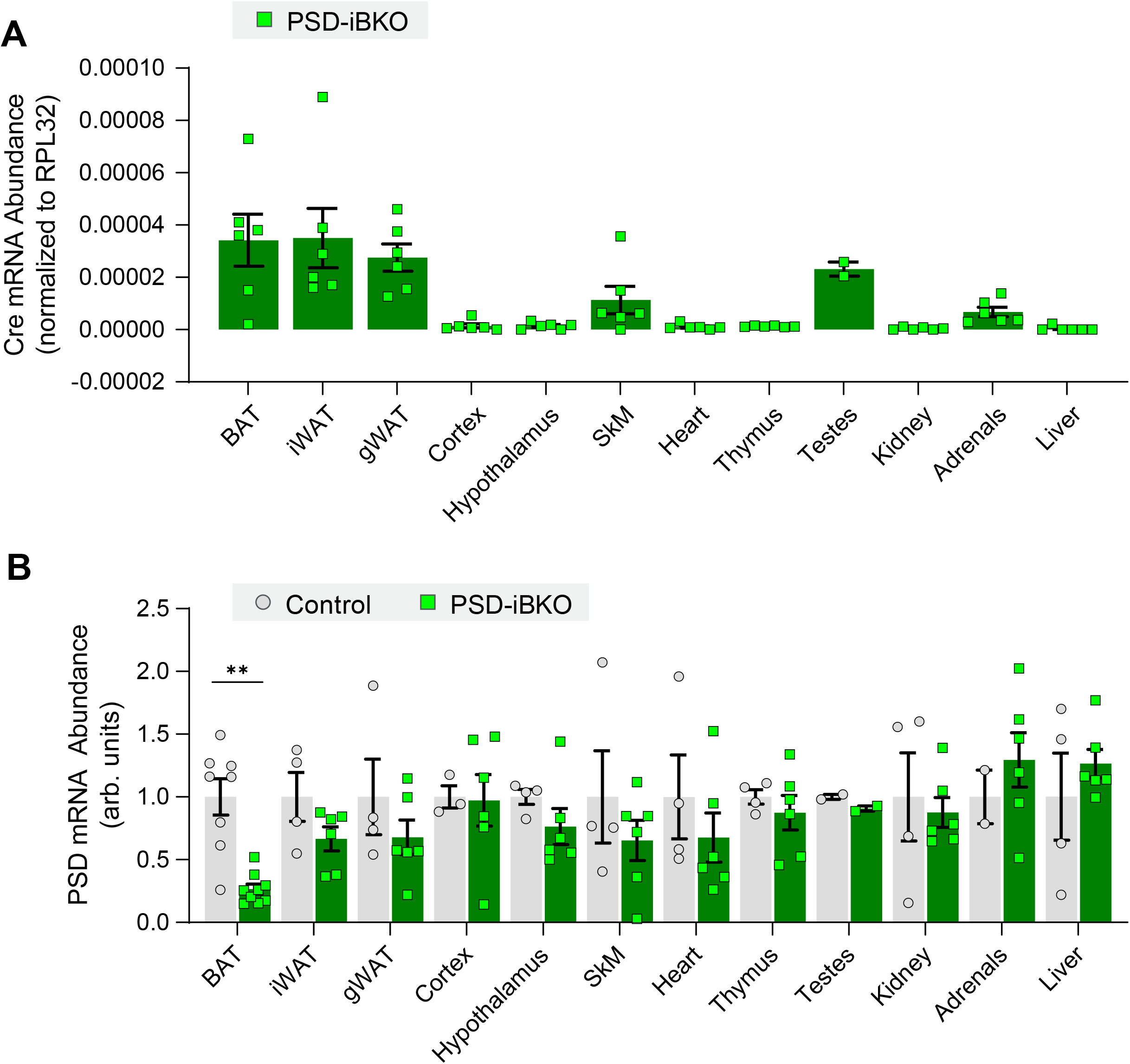
UCP1-CreERT2 expression does not result in decreased PSD expression outside of thermogenic adipose tissue. (A) Cre mRNA abundance in various tissues of PSD-iBKO mice. n=2-6/group. (B) PSD mRNA abundance in various tissues of control and PSD-iBKO mice. n=4-10/group. Data are presented as ± S.E.M. *Denotes of p-value of 0.05 or less.

**Supplemental Figure 6:**
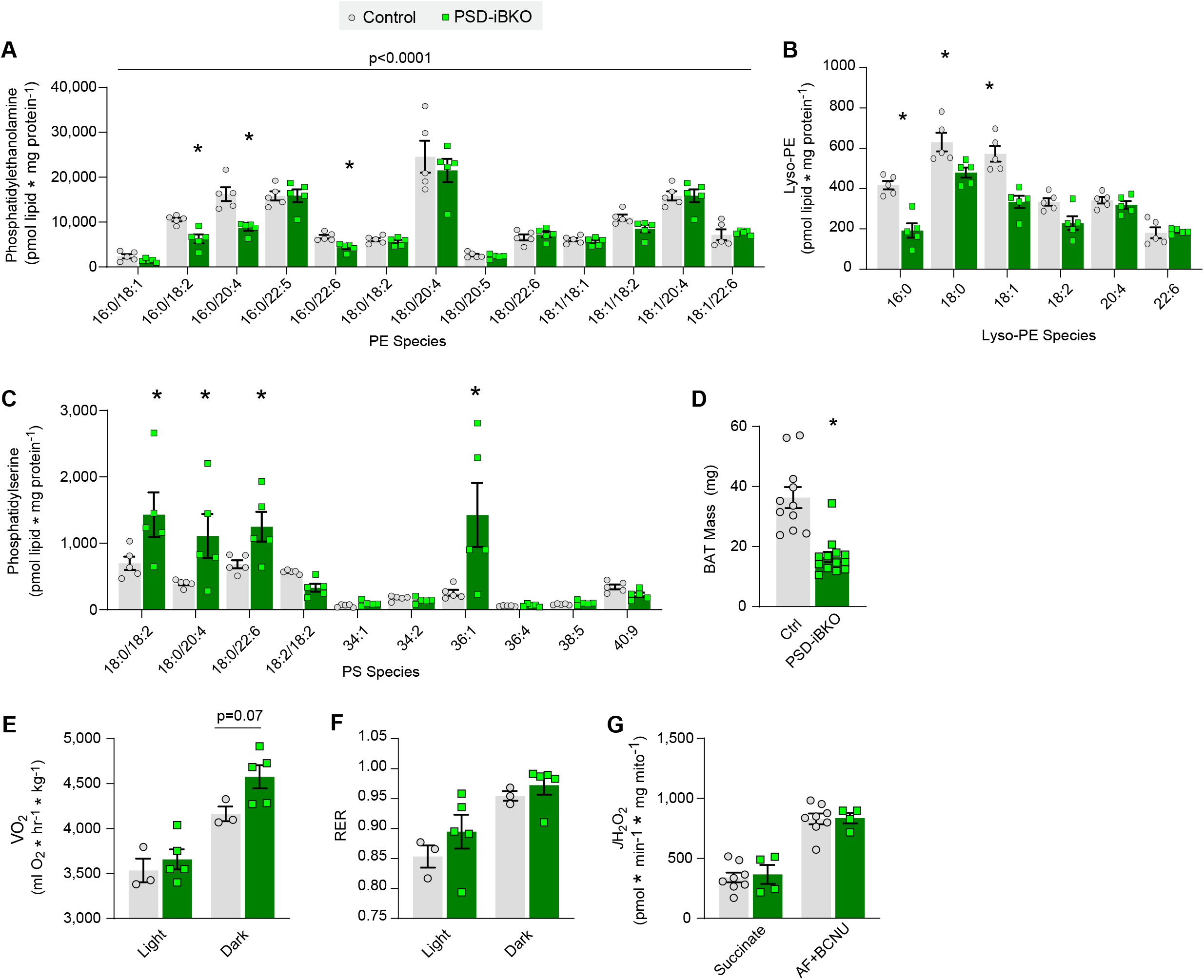
Additional data on PSD-iBKO mice. (A) Mitochondrial PE levels in BAT from control and PSD-iBKO mice, 4 weeks post-tamoxifen injection. n=7-13/group. (B) Mitochondrial lyso-PE levels, 2 weeks post-tamoxifen injection. n=5/group. (C) Mitochondrial PS levels, 2 weeks post-tamoxifen injection. n=5/group. (D) BAT mass. n=11-13/group. (E) Whole-body oxygen consumption in metabolic cage. n=3-5/group. (F) Respiratory exchange ratio (RER) in metabolic cage. n=3-5/group. (G) Mitochondrial H_2_O_2_ production in the presence of 10 mM succinate and antioxidant inhibitors auranofin (AF) and carmustine (BCNU). n=5/group. Data are presented as ± S.E.M. *Denotes of p-value of 0.05 or less.

**Supplemental Figure 7:**
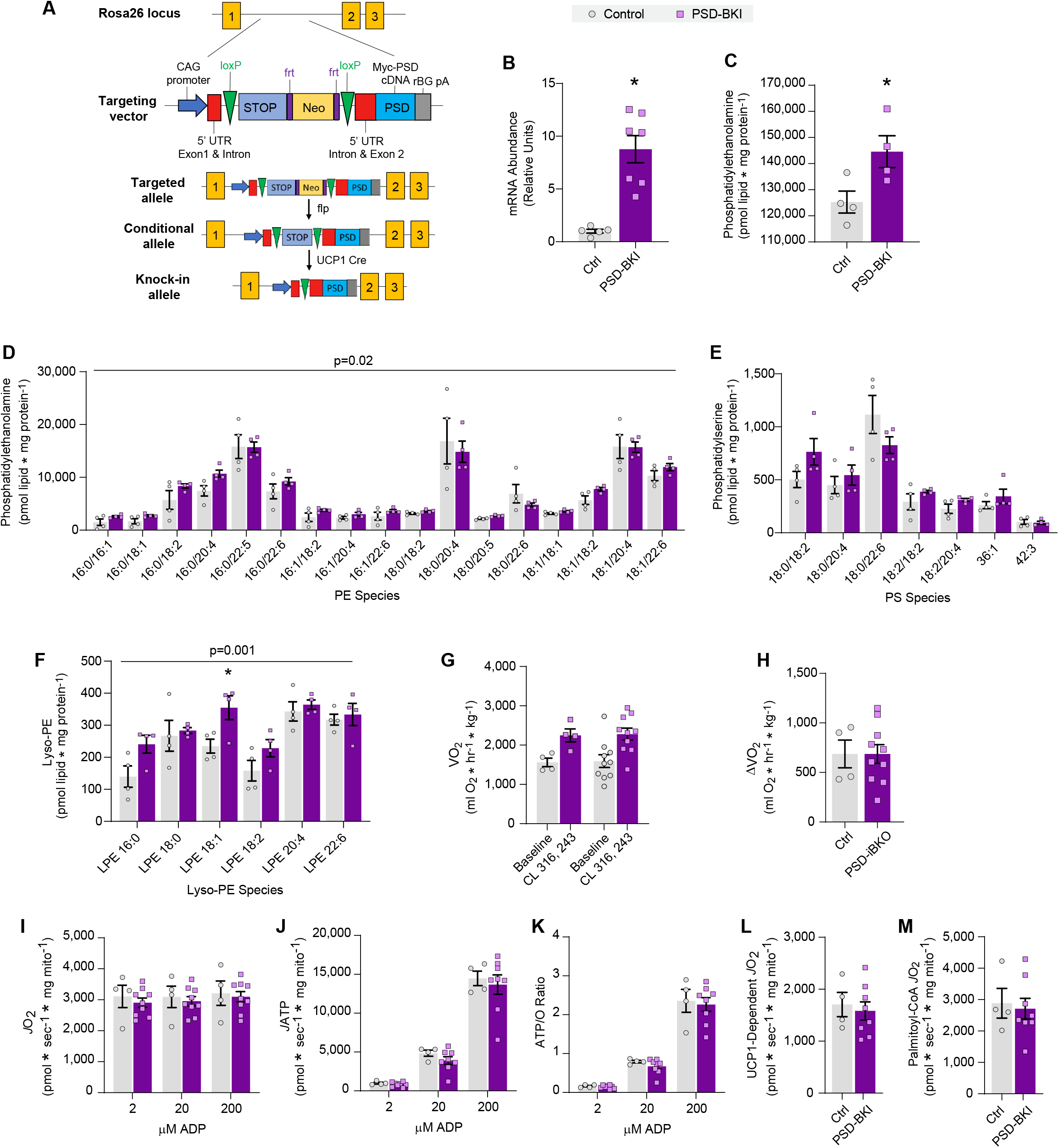
BAT-specific overexpression of PSD does not robustly influence mitochondrial PE, nor does it promote phenotypes in thermogenic capacity or UCP1-dependent respiration. (A) A schematic of the genetic strategy used to generate PSD-BKI mice. (B) PSD mRNA abundance in BAT from control and PSD-BKI mice. n=5-7/group. (C) Total mitochondrial PE in BAT from control and PSD-BKI mice. n=4/group. (D) Mitochondrial PE species in BAT. n=4-6/group. (E) Mitochondrial PS species in BAT. n=4-6/group. (F) Mitochondrial lyso-PE species in BAT. n=4-6/group. (G) Whole-body oxygen consumption before and after CL 316,243 administration. n=4-10/group. (H) Changes in whole-body oxygen consumption induced by CL 316,243 administration. n=4-10/group. (I) Mitochondrial O_2_ consumption in BAT from control and PSD-BKI mice, measured in the presence of 5 mM pyruvate, 0.2 mM malate, 5 mM glutamate, and 5 mM succinate, and 2, 20, and 200 µM ADP. n=4-8/group. (J) ATP production in the presence of 5 mM pyruvate, 0.2 mM malate, 5 mM glutamate, 5 mM succinate, and 2, 20, and 200 µM ADP. n=4-8/group. (K) Mitochondrial coupling efficiency (ATP/O ratio) of BAT mitochondria. n=4-8/group. (L-M) UCP1-dependent respiration in control and PSD-BKI mice stimulated by 5 mM pyruvate and 0.2 mM malate (L) or 0.2 mM malate, 5 mM carnitine, and 20 µM palmitoyl-CoA (M) and inhibited by 4 mM GDP. n=4-8/group. All phenotyping for control and PSD-BKI were done in mice housed in thermoneutrality for 4 weeks. At room temperature housing, PSD-BKI mice did not exhibit greater mitochondrial PE content compared to control mice. Data are presented as ± S.E.M. *Denotes of p-value of 0.05 or less.

## STAR METHODS

### KEY RESOURCES TABLE

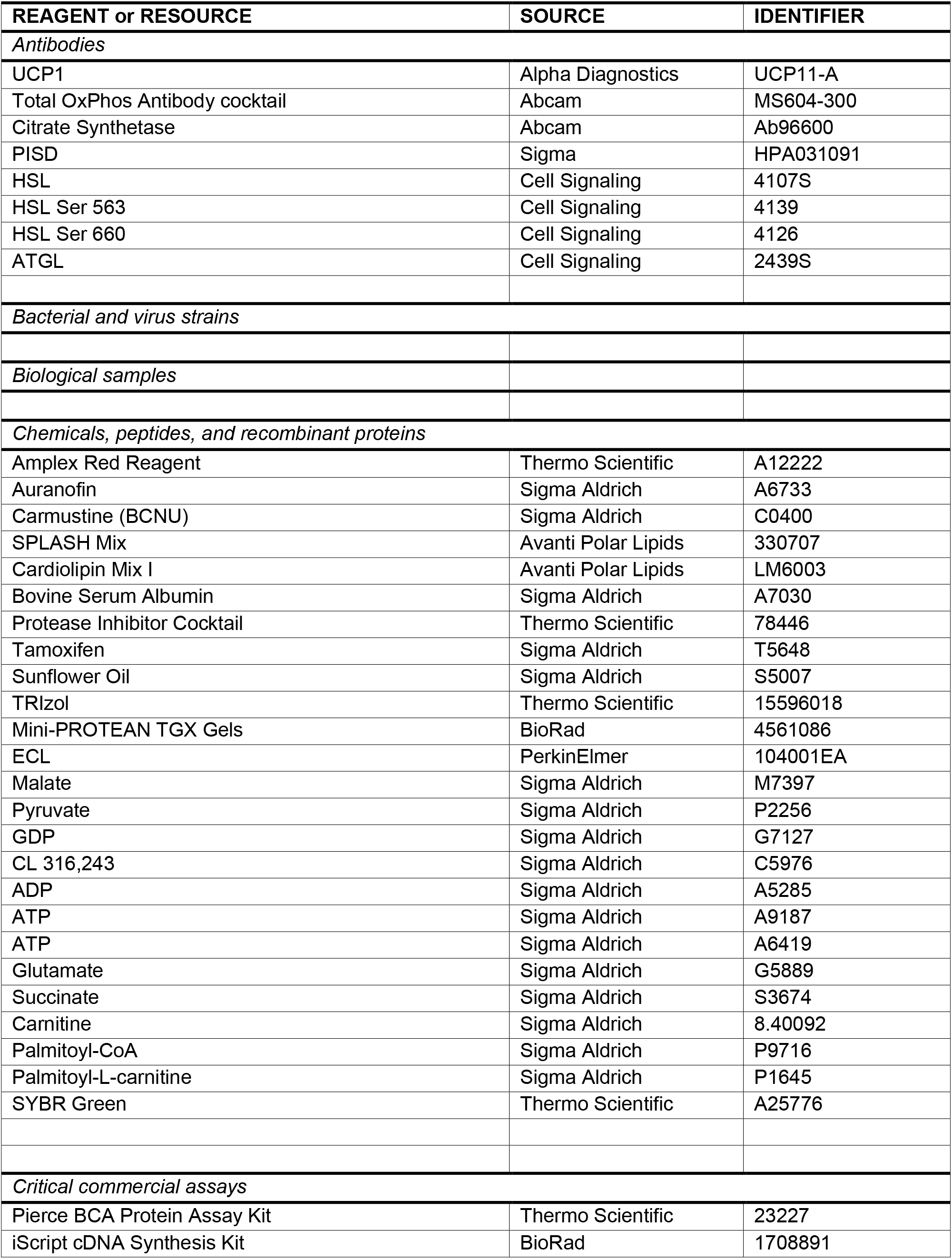
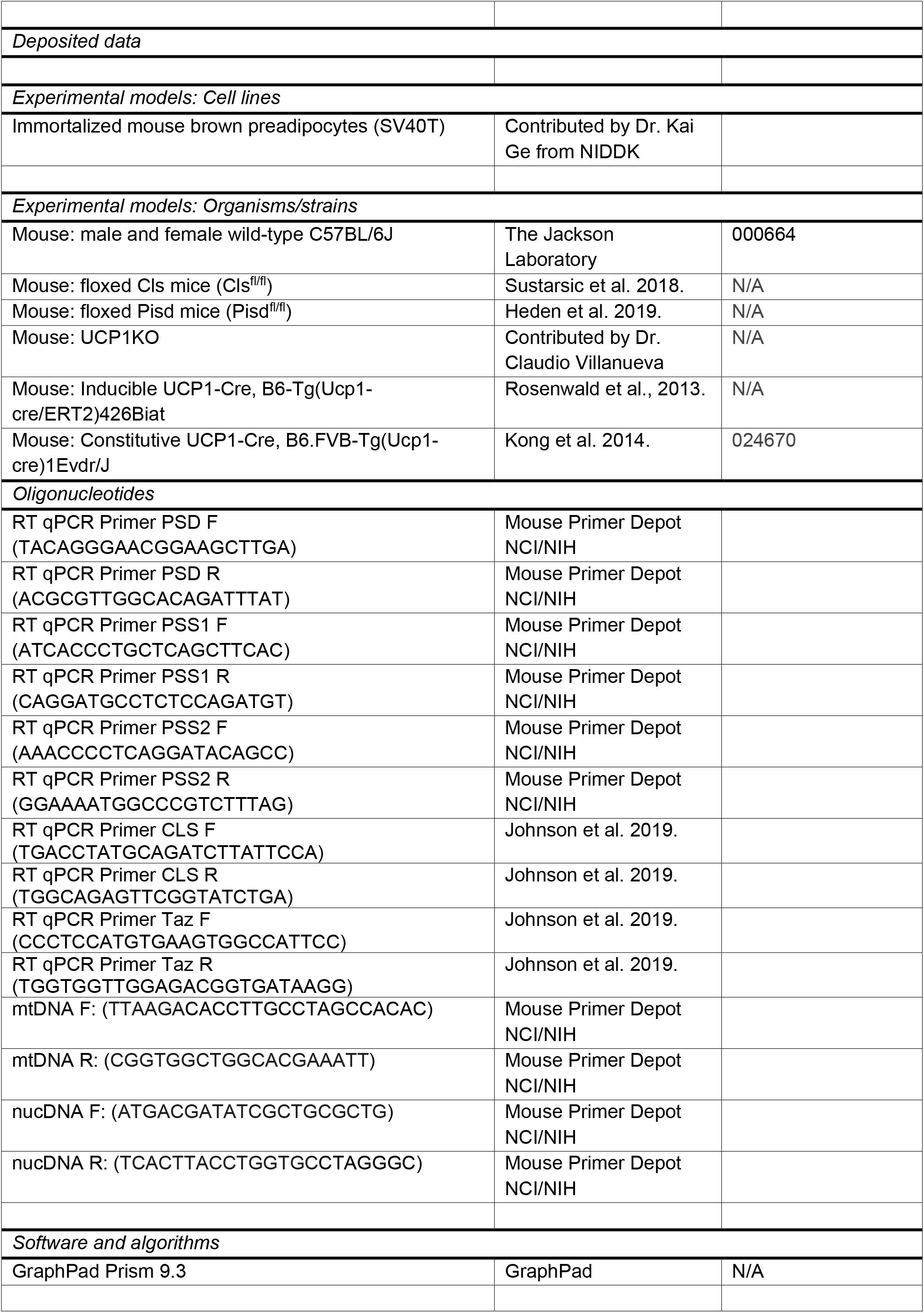
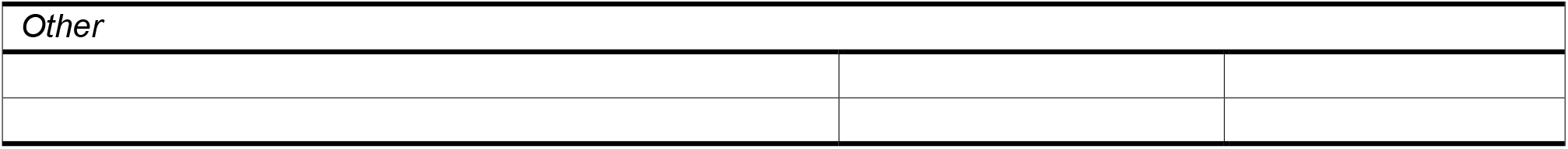

## RESOURCE AVAILABILITY

### Lead Contact

Please direct requests for additional information and resources and reagents to the lead contact for this manuscript, Dr. Katsuhiko Funai.

### Materials Availability

Plasmids utilized by this study are available from Sigma Aldrich. Mouse lines generated by this study may be available at personal request from the lead contact. No new reagents were created or used by this study.

### Data and Code Availability

The data generated by this study including all images, figures, and datasets, is available upon request to the lead contact, Dr. Katsuhiko Funai. Similarly, any additional information necessary to reanalyze datasets is also available upon request. No code was utilized in this study.

## EXPERIMENTAL MODEL AND SUBJECT DETAILS

### Genetically Modified Mice

The animal studies performed in this manuscript were approved by the University of Utah Institutional Animal Care and Use Committee. All mice utilized in this study were of the C57Bl/6J background. Unless otherwise noted, experiments were performed on mice 12 weeks of age that were housed at an ambient temperature of 22° C. They were fed a standard chow diet (Teklad 2020X). Transgenic animals were injected with 7.5 mg/kg of tamoxifen (Sigma-Aldrich T5648) dissolved in sunflower oil (Sigma-Aldrich S5007) for 5 days in a row 3 weeks (PSD-iBKO) or 4 weeks (CLS-iBKO) prior to sacrifice unless otherwise noted. Males and females were utilized for each experiment and no sex-dependent differences were observed. The mice were fasted for 4 hours prior to sacrifice using ketamine/xylazine.

#### Wildtype Mice

C57BL/6J mice stock # 000664 were obtained from The Jackson Laboratory.

#### CLS-iBKO

This study utilized tamoxifen-inducible BAT specific CLS knockout (CLS-iBKO) mice generated by Elahu Sustarsic and Dr. Zachary Gerhart-Hines (University of Copenhagen) (Sustarsic et al., 2018). This line was later mated to a UCP1-specific Cre-ERT2 driver (UCP1 Cre-ERT2^+/-^) generated by Dr. Christian Wolfrum (ETH Zürich) (Rosenwald et al., 2013). The resulting (CLS-cKO^+/+^, UCP1 Cre-ERT2^+/-^; designated as CLS-iBKO) and littermate control (CLS-cKO^+/+^, no Cre) mice were used in this study. Both groups were administered tamoxifen.

#### PSD-iBKO

Tamoxifen-inducible PSD knockout in BAT (PSD-iBKO) mice were generated for this study. The PSD conditional knockout (PSDcKO) mice were previously generated by the Funai lab (Heden et al., 2019). This line was then mated to the same UCP1-specific Cre-ERT2 driver (UCP1 Cre-ERT2^+/-^) from Dr. Christian Wolfrum’s Lab. Tamoxifen-injected PSD-iBKO (PSD-cKO^+/+^, UCP1 CreERT2^+/-^; designated as PSD-iBKO) and littermate control (PSD-cKO^+/+^, no Cre) mice were used in this study.

#### PSD-BKI

The BAT PSD knock-in (PSD-cKI^+/+^) mouse line was generated by inserting myc-tagged *Pisd* cDNA into a Rosa26 locus. The gene is preceded by a CAG promotor as well as a stop codon flanked by loxP sites to allow for tissue specific expression (lox-STOP-lox). The PSD-BKI line was created by mating PSD-cKI^+/+^ mice to mice with a constitutive UCP1-Cre driver (UCP1Cre^+/-^) obtained from Jackson Laboratories, stock # 024670 (Kong et al., 2014). This cross generated PSD-BKI (PSD-cKI^+/-^, UCP1Cre^+/-^; designated PSD-BKI) and littermate control (PSD-cKI^+/-^, no Cre) mice.

## METHOD DETAILS

### Mitochondrial Isolation

Mitochondrial isolation from BAT was performed as previously described (Johnson et al., 2020). BAT from freshly sacrificed mice was finely minced in mitochondrial isolation media (MIM, 300 mM sucrose, 10 mM HEPES, 1 mM EGTA) with 1 mg/mL BSA. The mixture was homogenized at 1000 rpm for 6-8 passes. Homogenates were centrifuged at 10,000 x g for 10 minutes. The supernatant was discarded and the samples were centrifuged at 200 x g for 5 minutes twice. Each time, the supernatant was collected (the pellet discarded) and transferred to a new tube. The supernatant was centrifuged once more at 10,000 x g for 10 minutes. The supernatant was discarded and the pellet was resuspended in MIM. Protein concentration was assessed using the Pierce BCA protein assay kit (Ref 23227 Thermo Scientific).

### Respirometry

Mitochondrial oxygen consumption was measured using Oroboros oxygraphs. Resorufin or NADPH production were measured using FluoroMax-4 (Horiba Scientific) for H_2_O_2_ and ATP assays respectively. Experiments were performed in buffer Z (105 mM K-MES, 30 mM KCL, 1 mM EGTA, 10 mM K_2_HPO_4_, 5 mM MgCl_2_-6H_2_O, 5 mg/mL BSA, pH 7.4) as previously described (Perry et al., 2011). UCP1-dependent respiration was stimulated by treating mitochondria with 0.5 mM malate and 5 mM pyruvate. UCP1 was inhibited using 4 mM guanosine diphosphate (GDP). ATP production was indirectly measured using an enzymatically coupled reaction that produced nicotinamide adenine dinucleotide phosphate (NADPH). NADPH was excited at 340 nm and fluorescence was measured at 460 nm every 2 seconds as previously described (Lark et al., 2016). ATP synthesis was stimulated by treating mitochondria with 0.5 mM malate, 5 mM pyruvate, 5 mM glutamate, and 5 mM succinate with 2 µM ADP, 20 µM ADP, and 200 µM adenine diphosphate (ADP). The ATP/O ratio was calculated by coupling the FluoroMax ATP assay with an identical assay performed in the Oroboros oxygraphs. H_2_O_2_ production in isolated mitochondria was stimulated using 10 mM succinate, followed by 1 µM auranofin and 100 µM carmustine (BCNU). Resorufin is produced at a 1:1 ratio with H_2_O_2_ in the presence of Amplex Red, superoxide dismutase, and horse radish peroxidase. It was excited at 550 nm and fluorescence was measured at 585 nm.

### Lipidomics

Mitochondrial phospholipids were extracted from isolated mitochondria using a modified Matyash lipid extraction protocol (Matyash et al., 2008). Phospholipid internal standards (SPLASH Mix Avanti Polar Lipids 330707 and Cardiolipin Mix I Avanti Polar Lipids LM6003) and 50 µg of protein from isolated mitochondria were added to ice cold 3:10 methanol:methyl-tert-butyl-ether. Samples were vortexed and sonicated for 1 minute before being incubated on ice for 15 minutes. During this time, samples were vortexed every 5 minutes. H_2_O was added and the samples were again incubated on ice for 15 minutes with vortexing taking place every 5 minutes. The samples were centrifuged at 15,000 x g for 10 minutes. The supernatant was harvested and the solvent evaporated using a SpeedVac set to 37 °C for 1 hr. The lipid pellet was resuspended using a 9:1 methanol:toluene mixture. Phospholipid analysis was conducted using liquid chromatography-mass spectrometry (LC-MS) on an Agilent 6530 UPLC-QTOF mass spectrometer.

### Metabolic Phenotyping

Whole body VO_2_ was measured using a Comprehensive Lab Animal Monitoring System (Columbus Instruments). Cold-tolerance testing was conducted at 4 °C. Core body temperature was measured using a temperature-sensitive transponder injected into dorsal subcutaneous adipose tissue (Bio Medic Data Systems, IPTT 300). The transponder was placed in the mice one week prior to cold tolerance testing. Transponder readings were assessed using a Reader-Programmer (Bio medic Data Systems, DAS 8007). Mice were single-housed with access to food, water, and bedding during cold tolerance testing. Mice were euthanized after 4 hours of fasting.

### Electron Microscopy

BAT was isolated from mice, minced with scissors, and fixed in ice-cold 2.5% glutaraldehyde. After incubating for 48 hours at 4 °C, the tissue was delivered to the University of Utah Electron Microscopy Core for post-fixation dehydration and embedding. The embedded tissues were sectioned and stained using uranyl acetate. The tissues were then imaged with a JEOL JEM1400 Plus transmission electron microscope and acquired using a Soft Imaging Systems MegaView III CCD camera.

### Gene Expression

BAT stored at -80° C was thawed and placed in 1 mL of TRIzol. The tissue was homogenized and spun down at 10,000 x g for 10 minutes. The lipid layer was discarded and 200 µL of chloroform was added to the tube. The tube was mixed by inverting it 10 times. After 2 minutes, samples were centrifuged at 13,000 x g for 15 minutes at 4 °C. The aqueous supernatant was collected and placed in a new tube containing 500 uL of 100% isopropyl alcohol. The mixture was inverted a few times and incubated at room temperature for 10 min. The samples were spun down at 13,000 x g for 10 minutes at 4 °C. The supernatant was carefully aspirated such that the pellet was not disturbed. The pellet was washed with 75% ethyl alcohol and spun down at 5,500 x g for 5 min. The supernatant was aspirated and the pellet was air dried for 10 minutes before being resuspended in Tris-EDTA buffer. A cDNA library was generated by reverse transcribing the RNA using the iScript cDNA Synthesis Kit (Bio-Rad). For quantitative PCR, cDNA was combined with SYBR Green (Thermo Fisher Scientific) and gene specific primers and then placed in a 384 well plate. Gene expression was analyzed using a QuantStudio 12K Flex (Life Technologies).

### Histological Analysis

BAT was fixed by placing the tissue in a 4% paraformaldehyde PBS solution for 48 hours. The tissue was then placed in 70% ethanol for another 48 hours. The tissue was embedded in paraffin, cut into approximately 5 µm pieces, and stained using hematoxylin and eosin or Masson’s trichrome blue. The sectioned tissues were then imaged using an Axio Scan.Z1 (Zeiss).

### Protein Analysis

For whole cell analysis, BAT stored at -80 °C was thawed and placed in ice cold homogenization buffer (150 mM NaCl, 50 mM Tris-HCl, 5 mM EDTA, 1% Triton X-100, 0.1% SDS, 0.1% sodium deoxycholate, and 1% protease and phosphatase inhibitor cocktail added immediately prior to use) (Ref 78446 Thermo Scientific). The sample was homogenized and centrifuged at 12,000 x g for 5 minutes. The concentration of protein was determined using a BCA (details included above). For analyzing proteins in isolated mitochondria, mitochondria were first isolated from BAT using the protocol outlined above. From this point forward, identical protocols were followed. Equal quantities of protein were mixed with Laemmli sample buffer and loaded into a gradient SDS-PAGE gel (Ref 4561086 Bio-Rad). The proteins were transferred from the gel onto nitrocellulose membranes. Membranes were blocked using 5% BSA in TBST for 1 hour before being treated with primary antibodies overnight at 4 °C. The membranes were washed 5 times using TBST and placed in secondary antibodies for 1 hour. The membranes were washed 5 times in TBST and twice in TBS before imaging. Immediately prior to imaging, ECL (Ref 104001EA PerkinElmer) was pipetted onto the membrane. Membranes were imaged using a FluorChem E imager (ProteinSimple).

### Isolation of BAT mitoplasts

5 week old mice were sacrificed (CO_2_ asphyxiation) followed by cervical dislocation. Interscapular BAT was manually separated, and mitochondria were isolated as described in (Balderas et al., 2022). Tissue was disrupted with a Potter-Elvehjem homogenizer, and a crude mitochondrial fraction isolated by differential centrifugation. Mitoplasts were generated from BAT mitochondria using a French press set at 1200 psi to mechanically break the OMM. For recording, mitoplasts were aliquoted into a divalent-free KCl (DVF KCl) solution containing 150 mM KCl, 10 mM HEPES, and 1 mM EGTA (pH 7.2 with KOH) and plated on 5 mm coverslips coated with 0.1% gelatin.

### Whole mitoplast electrophysiology

Mitoplasts (3–5 µm) had typical membrane capacitances of 0.5–1.0 pF. 5-10 GΩ seals were formed in DVF KCl, where a fast voltage step of 250–600 mV was applied to break-in. Entering the whole-mitoplast configuration was monitored by changes in the amplitude of the capacitance transients. Mitoplasts were interrogated every 4 seconds with a ramp protocol from - 160 to +80 mV, at a holding potential of 0 mV. Recording borosilicate pipettes (15-25 MΩ resistance) were filled with 150 mM Tetraethylammonium hydroxide, 1.5 mM EGTA, 1.0 mM magnesium gluconate, 150 mM HEPES, and 2 mM Tris-Cl (pH adjusted to 7.0 with D-gluconic acid, tonicity adjusted to 325-350 mmol/kg with sucrose). After establishing whole-mitoplast configuration in KCl DVF solution, the bath solution was changed to a proton current recording solution containing 150 mM HEPES, 1 mM EGTA, and 0.5 mM MgCl_2_ (pH adjusted to 7.0 with Tris-base, tonicity adjusted to 300 mmol/kg with sucrose). Data acquisition and analysis were performed using PClamp 10 (Molecular Devices) and Origin 7.5 (Origin Lab). Electrophysiological data were acquired at 10 kHz and filtered at 0.5-1 kHz. Images were prepared on Adobe Illustrator. For presentation purposes of exemplars, these have been subject to the Simplify filter to reduce file size without changing shape, and fast capacitance transients have been removed.

## QUANTIFICATION AND STATISTICAL ANALYSIS

Data were analyzed using GraphPad Prism 9.3 software. The value of n for each experiment is noted in the figure legends and corresponds to data obtained from an individual mouse or batch of cells. All data are presented as means ±SEM. Significance was set at p < 0.05. Data with only 2 groups was analyzed using two-tailed Student’s *t* test. Data with more than 2 groups was analyzed using one-way ANOVA.

